# *GOParsimony*: parsimonious saltation across the Gene Ontology digraph for efficient functional annotation

**DOI:** 10.64898/2026.11.01.763467

**Authors:** Paul M. Harrison

## Abstract

**Background:** The Gene Ontology (GO) is standardly probed by biological researchers keen to discern the functional tendencies of their gene/protein data sets. However, lists of GO terms are often ontically overwhelming, and need to be filtered to remove redundancy. Methods for such filtering either employ predefined lists of terms that may not actually be relevant to a specific gene/protein set, or apply arbitrary thresholds, such as for semantic distance (*i*.*e*., GO term dissimilarity).

**Methods:** Here, I introduce the tool GOParsimony, which is designed to avoid such issues and to pick out the most parsimonious (*i*.*e*., simplest, most concise) explanation for the functional tendencies of any set of proteins/genes. It achieves this by saltating across the GO digraph, assigning and updating cascades of term precedence until convergence. The set of GO terms assigned precedence = 1 is the most parsimonious set. Hierarchical clustering with other GO terms with higher precedence levels can be investigated for clues about functional modules.

**Results:** GO term precedence graphs and two-dimensional heatmaps of GO terms and genes/proteins are output, along with other plots that can be cross-referenced for functional insights. In test analyses for five example data sets from budding yeast and human, GOParsimony consistently produces reduced GO term lists that have greater significance and low or intermediate coverage of the space of significant GO terms, when compared to other distributed methods in default mode. GOParsimony can be obtained at: https://github.com/pmharrison/GOParsimony.

## Introduction

Hypothesis generation about protein or gene function is the primary goal of cell and molecular biology investigations. The Gene Ontology (GO) is an ever-growing resource of functional classifications and annotations that is routinely mined for such hypotheses (Eden et al. 2009; Ge et al. 2020; Gene Ontology 2026). During this data mining, GO terms are taken from the GO digraph to label onto proteins/genes. However, often this mining is ontically overwhelming, *i*.*e*., there is too much detail in the functional annotations, and it is not clear which GO terms take precedence over others. GO provides a smaller pre-determined set of functional terms called GOSlims to reduce the complexity of term mapping (Gene Ontology 2026). However, such lists might not be relevant to specific gene/protein data sets and could miss key features.

Other approaches to reducing the complexity of GO annotation involve the application of semantic distance thresholds for GO terms (i.e., GO term dissimilarity), such as in the tools Revigo and GO-Figure! (Reijnders & Waterhouse 2021; Schlicker et al. 2006; Supek et al. 2011). Drawbacks of such calculations are the reliance on arbitrary distance thresholds and the variety of possible distance definitions. Other approaches such as the TopGO elim/weight and parentchild algorithms, rely on the comparisons of the protein/gene annotations for relative parent and child GO digraph terms to penalize the significance of parent nodes; such continual re-weightings might excessively favour outputs to more specific GO terms, thus producing some false positives rankings of terms (Alexa & Rahnefuhrer 2026; Alexa et al. 2006; Grossmann et al. 2007).

Here, to avoid such issues, the GOParsimony tool has been developed. It picks out the most parsimonious (*i*.*e*., simplest, most concise) explanation for the functional tendencies of any set of proteins/genes. It achieves this by saltating across the GO digraph, assigning and updating cascades of term precedence until convergence. The set of GO terms assigned precedence = 1 is the most parsimonious set. Hierarchical clusterings with other GO terms with higher precedence levels can be investigated for clues about functional modules.

## Methods

### Algorithm

GOParsimony is an algorithm designed to derive the most parsimonious explanation (*i*.*e*., the simplest, most concise) for the functional emphases of a gene/protein data set.

GOParsimony proceeds through four stages (Figure 1A). It needs to be fed a *background* set of Gene Ontology (GO) term annotations, which might be for the whole proteome, or the whole set of genes tested in an expression experiment. This is input in GO GAF format (Gene Association File, *.gaf) or a simpler two-column format (gene/protein ID + GO term). Secondly, a *sample* set of IDs or accessions is input (*i*.*e*., for the sample of proteins/genes the user is studying).

**Figure 1.**
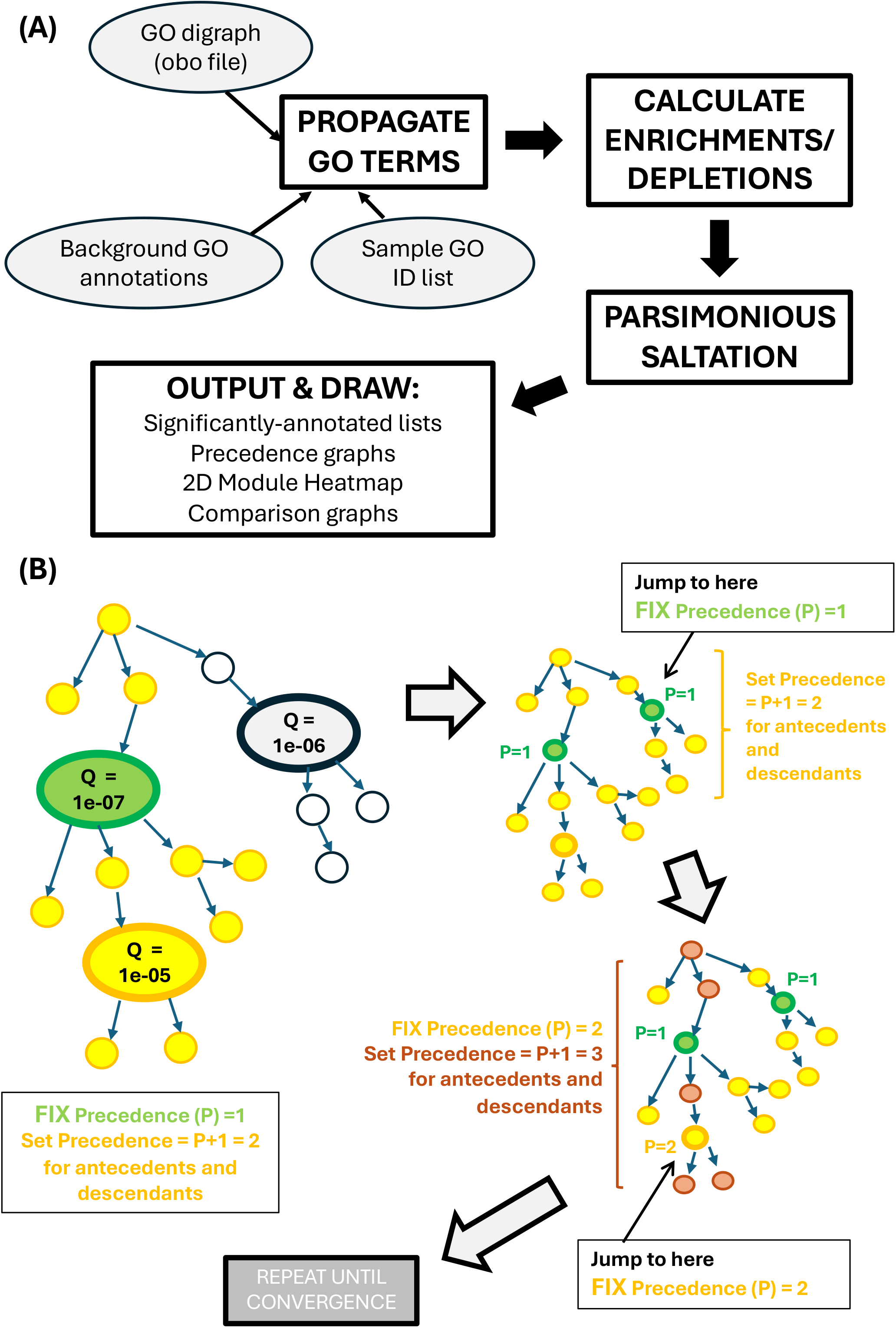
GOParsimony overview. (A) The four stages of the pipeline as detailed in Algorithm. (B) The parsimonious saltation algorithm for assigning precedence levels to terms. The detailed algorithm is explained in the Algorithm section. A schematic of part of a GO digraph of significant terms is depicted. The three terms labelled with Q-values (i.e., P-values corrected by the Benjamini-Hochberg procedure) are processed. Green colour indicates precedence level = 1, yellow precedence level = 2, and brown precedence level = 3. Three steps of the saltation (‘jumping’) and precedence level setting/fixing are shown.

In the first stage, more general GO overterms that are missing are propagated onto the input sample and background set from consulting the GO term digraph. This digraph is in *.obo file format, and can be provided by the user, or otherwise the default digraph file bundled with the package is input. The resulting expanded lists of annotations can be optionally output.

In the second stage, enrichments and depletions of GO terms in the sample relative to the background are calculated and assigned a significance using hypergeometric probability. A Benjamini-Hochberg correction for multiple hypotheses is applied (a Bonferroni correction is also possible) (Benjamini & Hochberg 1995). Depletions are calculated by default but are not output.

The third stage is *parsimonious saltation* (Figure 1A-B). This is a parsimonious procedure for reducing redundancy in the annotations. In the toy digraph (Figure 1B), three significant GO terms and their antecedent (parent) and descendant (child) terms are assessed. For the most significant one, the *precedence* level (P) is fixed at = 1, and all of its antecedent and descendant terms are set to precedence level 2 (= P+1). We then saltate (jump) to the next most significant term, fix its precedence level = 1 and set its antecedents and descendants to precedence level 2. Then finally, we jump to the third term fix its precedence level = 2, and set = 3 the precedence of any of its antecedent and descendant terms that do not have fixed precedence. This algorithm then repeats until convergence for all remaining GO terms. The *most parsimonious* (MP) set of terms fixed to precedence level 1 is the core list reduced for annotation redundancy, however the higher precedence levels need to be probed to gain insight into possible functional modules, as discussed below in *Examples*.

Finally, plots and tables are output. These include the lists of significantly annotated proteins/genes, *precedence graphs* that highlight the patterns of precedence (both full-size and concise formats), and two-dimensional heatmaps of use for picking out potential functional modules (in these, Jaccard distances derived from GO-term or protein/gene list membership overlaps are used to generate the hierarchical trees). Also, there are two sorts of *comparison graph*: (i) a bar chart which indicates whether further functional detail is added from consideration of higher precedence levels, and (ii) a graphical comparison of the precedence=1 set with GOslims annotation (if a GOslims file is specified) (Gene Ontology 2026), and a input file of highlighted GO terms if such a file is provided by the user (this could be from another method to assess GO term redundancy). Precedence is abbreviated to *Prec* in all graphical outputs.

### Implementation & distribution

The package is implemented in C (*GOParsimony*.*c*, which compiles to *GOParsimony*), Python (*precedence-graphs*.*py*), and shell script (*run-GOParsimony*.*sh*). It runs on Mac OSX or Linux. Installation and compilation are explained in the bundled README.txt file. Google Gemini (google.gemini.com) was used to edit and revise code. Compilation and running were tested in Mac OSX Tahoe 26.5, and in Ubuntu 22.04.5 LTS within a Docker wrapper, with a variety of options and input files. The complete GOParsimony is distributed at https://github.com/pmharrison/GOParsimony, with example inputs and outputs. The DOI for the Zenodo archive of GOParsimony is DOI 10.5281/zenodo.20543570. Further Supplementary Info (namely textual outputs for each of the five examples) have been deposited at Zenodo: https://zenodo.org/records/19949239.

### Options

When run with the script *run-GOParsimony*.*sh*, the package has the following options:

–b background annotations file (required)

–s sample ID/accession list (required)

–o GO obo file (if not specified the bundled default is used)

–m a GOslims file in *.obo format

–l ‘ell’, a GO term highlights file (just a single-column list)

–p an output file prefix

–x maximum precedence level to use in graphical outputs (default=5)

–e to output annotation lists expanded to include missing overterms

The GOParsimony C program has additional options that are not available when running the whole package: for querying a GO term from the ontology file (–q option), for setting a significance or false discovery rate threshold other than 0.05 (–d), for a concise textual output (– c), for enabling analysis and output of depletions (–D), and instead of the default Benjamini-Hochberg procedure, specifying Bonferroni correction (–B). Significant GO depletions tend to be more general terms, and as such they might indicate general deficiencies in the experimental characterization of proteins, such as insufficient protein-binding or localization data. Also, Bonferroni corrections tend to produce shorter lists of more general GO terms.

## Results

### Usage

GOParsimony is run on the command line. Here are some examples of its usage:

1. .*/run-GOParsimony*.*sh –o my-go*.*obo –b spleen-background*.*txt –s spleen-sample-IDs*.*txt –p spleen-expression* Here, the user specified its own obo-format GO digraph file my-go.obo and is analyzing a sample file of IDs called spleen-sample-IDs.txt. The background file is in two-column text format (gene/protein ID + GO term). The default maximum precedence level = 5 will be used for drawing graphs and plots, and output files will be prefixed with ‘spleen-expression’.
2. .*/run-GOParsimony*.*sh –b whale-proteome*.*gaf –s whale-prions-IDs*.*txt –m goslims*.*obo –x 3* In this run, the user is processing the background whale proteome annotations input in GO GAF format. A GOslims file is specified (this will appear in the graphical outputs and the textual tables). Maximum precedence = 3 is requested, and the outputs get no user prefix.
3. .*/run-GOParsimony*.*sh –b whale-proteome*.*gaf –s whale-prions-IDs*.*txt –m goslims*.*obo –x 3 –l whale-revigo-highlights*.*txt –e*

Here, the whale data is being analyzed again, but –e is used to output the expanded lists of GO term annotations for the sample and background input files, and a GO term highlights file is specified (–l, ‘ell’). Here the user is highlighting terms picked out by the Revigo website (Supek et al. 2011).

Gene Ontology digraph files in OBO format can be downloaded from https://geneontology.org/docs/download-ontology/. GO annotations can be obtained from https://geneontology.org/docs/download-go-annotations/ or from https://ftp.ebi.ac.uk/pub/databases/GO/goa/proteomes/ (suffixed *.goa), or from NCBI Refseq. Instructions to obtain them are at https://geneontology.org/docs/download-go-annotations/.

### Test examples

Five examples of GOParsimony application are discussed, three for proteins/genes sets from budding yeast (*Saccharomyces cerevisiae*), and two from human. These two organisms were picked since the Gene Ontology annotations for them are amongst the most detailed and comprehensive. The examples are used to describe the graphical output formats and their interpretations. These are collated in the five Supplementary Files (one for each example) provided along with this article, and the Figures therein are lettered A through F. The complete remaining textual input and output data for these examples are available at: https://zenodo.org/records/19949239.

#### (1) Cluster of budding yeast proteins with AQ-rich intrinsic disorder

Previously, compositionally-biased regions associated with intrinsic disorder (termed ID-CBRs) were clustered non-paralogously and discovered to have hundreds of functional associations (Harrison 2024a; Harrison 2024b). Here, we have picked out a small alanine/glutamine-rich (AQ-rich) cluster of nine proteins to pass through GOParsimony (Suppl. File 1). The *precedence graph* (Suppl. File 1, Figure A) indicates an MP set of five terms (Precedence = 1, deepest blue tint and bold text), that does not correspond with the GOslims annotations (magenta outline), or the more numerous list of terms obtained from default Revigo (Supek et al. 2011). Only the terms that are significant after the Benjamini-Hochberg correction for multiple hypotheses are shown, with dotted arrows for connections via undrawn non-significant terms. The *concise precedence graph* output is the same as the full precedence graph format, except that all the terms with precedence > than maximum precedence are omitted, and the arrows spanning their omission are dotted (Suppl. File 1, Figure B). The first *comparison graph* shows the lack of overlap between the three reduced sets of terms, with the MP precedence=1 set having the highest average –log(P-value), meaning the terms are on average more significant (Suppl. File 1, Figure E). The second *comparison graph* indicates that there is not much more detail to add in from considering higher precedence levels, since these additional levels tend to be parent terms (Suppl. File 1, Figure D). Finally, the 2D heatmap (Suppl. File 1, Figure C) indicates potential functional modules, *e*.*g*., P19659, P40356, P14922 and P22579 cluster together and are associated with the precedence=1 terms *transcription coactivator activity* and *response to abiotic stimulus*.

#### (2) EK-rich budding yeast proteins with low ‘blockiness’

In a recent study, glutamate/lysine-rich (EK-rich) ID-CBRs in *S. cerevisiae* were assessed for ‘blockiness’, *i*.*e*., the degree of segregation of residue types (Harrison 2024b; Harrison 2025). This is a set of 74 low-blockiness EK-rich ID-CBRs (Harrison 2024b) (Suppl. File 2). Although these proteins share the precedence=1 term *nuclear lumen* with the high-blockiness EK-rich set below, the higher precedence-level term patterns linked to *nuclear lumen* are different and generate a distinct heatmap. There is a notable set of 15 proteins largely linked to *cytoskeleton organization, response to stress, regulation of organelle organization* and *cell cycle process*. Also, the GOSlims and MP precedence=1 sets share no overlap, despite both covering ∼18% of the significant GO-term space.

#### (3) EK-rich budding yeast proteins with high ‘blockiness’

A list of 87 proteins that bear high-blockiness ID-CBRs was derived in the same study that example set (2) is from, and passed through GOParsimony (Suppl. File 3). The comparison graphs indicate that further functional clustering detail is gained from considering higher precedence levels, and the MP precedence=1 set has the highest mean term significances (highest mean –log[P-value]), compared to GOSlims and the set derived from default Revigo (Gene Ontology 2026; Supek et al. 2011). In the 2D heatmap, a cluster of twenty proteins 16 of which are labelled with the term *nucleolus* (a precedence level 3 term) stand out, as well as a smaller cluster linked to protein-DNA complex assembly (also precedence level 3). This demonstrates how these heatmaps might be useful for generating hypotheses about functional modules.

#### (4) Human intrinsically-disordered proteins associated with cancer (hcIDPs)

A data set of 143 human proteins annotated as containing experimental or predicted intrinsic disorder and associated with cancer by UniProt curators was derived and fed into the GOParsimony pipeline (UniProt 2023) (Suppl. File 4). The precedence=1 set here includes more specific GO terms such as *glucose homeostasis, establishment of protein localization to telomere, platelet activating factor receptor activity, glutamatergic synapse*, and more general terms such as *reproductive process, chromosome, nucleus* and *anatomical structure development*. The precedence=1 term *protein-containing complex* notably has a diverse array of precedence=2 children, including *beta-catenin destruction complex, CSF1-CSF1R complex, protein-DNA complex, transcription regulator complex, receptor complex*, and *Wnt signalosome*. The *comparison graphs* indicate that a lot of extra child detail is gained from considering higher precedences ≤3, and that the GOParsimony precedence=1 set is only 2% of the total list of significant GO terms, and has much greater average significance compared to GOSlims or default Revigo. There are many clusters on the 2D heatmap that might be informative for further functional hypotheses, *e*.*g*., a cluster of four proteins associated with *glucose homeostasis* and *DNA-binding transcription factor activity, RNA polymerase II-specific*.

#### (5) Genes up-regulated in human lymphoblasts cells treated with ionizing radiation

This is the data set used as a supplementary example in ref. (Ge et al. 2020) (Suppl. File 5).

As for the hcIDPs, the most parsimonious precedence=1 set is a mix of narrow and broad terms; also, we see the same phenomenon of a more general term labelled precedence=1 connected to diverse precedence = 2 or = 3 children. Here, in the concise precedence graph, *protein binding* (precedence=1), splits into 16 diverse precedence=3 binding types (including *signaling receptor binding, transcription factor binding, identical protein binding*, and *cyclin binding*). The trends in the *comparison graphs* are similar to those for the hcIDPs.

For the latter two human data set analyses, the maximum precedence level was set at ≤3, however, the graphs/plots are still quite large, and might only be suitable in publications as supplementary files. Another alternative is to extract ‘zoom-ins’ of favourite plot regions.

### Method Comparisons

The annotation reductions of several methods were compared for each of the five examples (Figure 2). Further detail for these methods appears in the *Background* section above. Jaccard distances derived from the overlaps between these lists of GO terms were employed to generate hierarchical trees. The data sets are:

**Figure 2.**
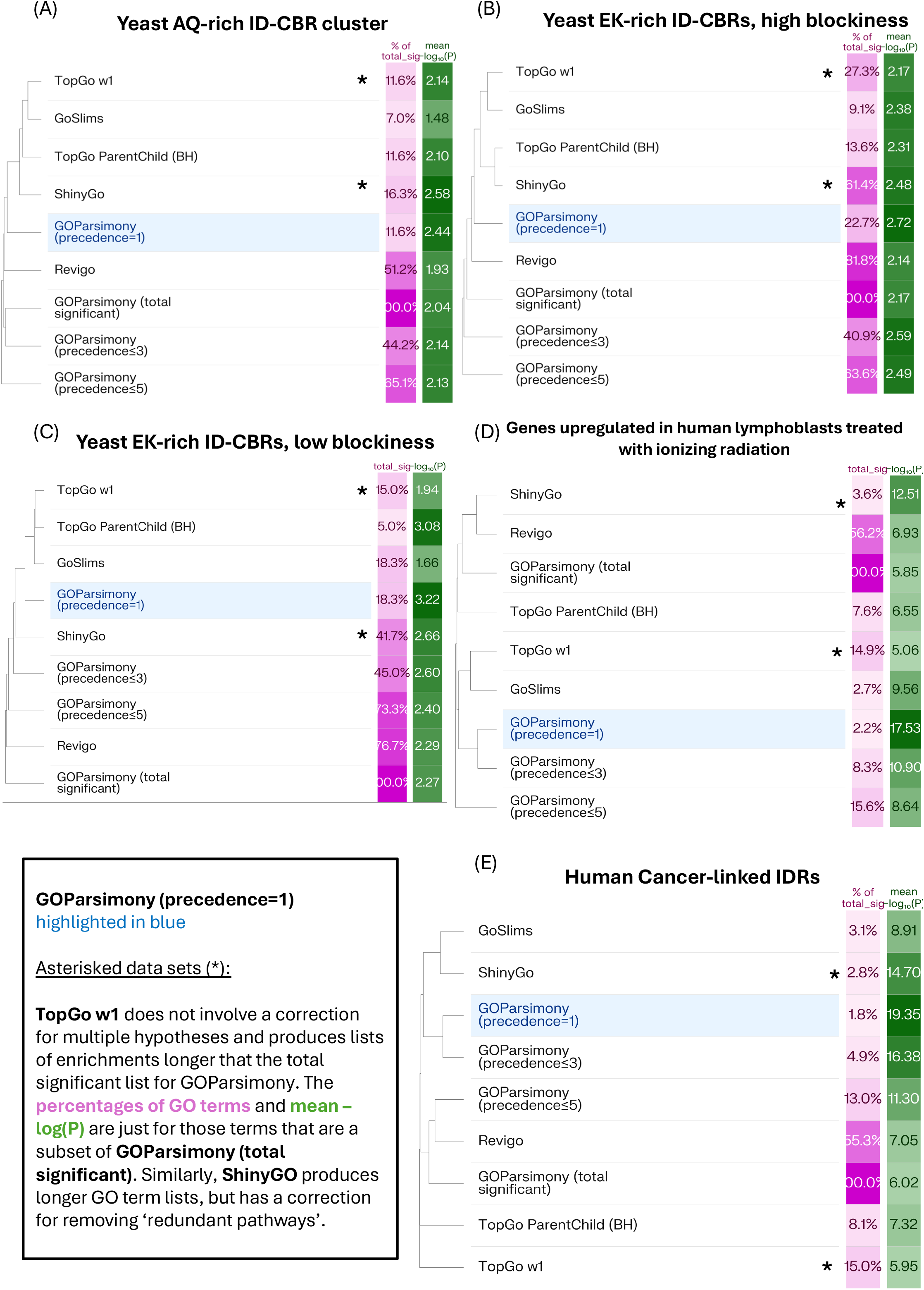
A comparison of different methods for processing GO terms for redundancy, for the five example data sets. The Revigo and ShinyGO annotations are made using default parameters, but with the P-value setting for term associations. A Benjamini-Hochberg correction is applied to the TopGO ParentChild algorithm , and to all other data sets except ShinyGO and TopGo w1. The percentage of total significant terms in the sets is colour-coded on a magenta scale, and the mean -log(P-value) is colour-coded on a green scale. The methods are hierarchically clustered using Jaccard distance calculated from the overlaps of the lists of significant GO terms produced by each one. This Figure was drawn by submitting these lists of GO terms to Claude Sonnet 4.8 (http://claude.ai) with appropriate queries.

i. the total list of significant GO terms (*i*.*e*., after correction for multiple hypotheses with the Benjamini-Hochberg method);
ii. the MP (most parsimonious) subset of the total significant list for GOParsimony (precedence=1);
iii. the subset for GOSlims (Gene Ontology 2026);
iv. the subset for Revigo (run with defaults) (Supek et al. 2011);
v. the subset after applying TopGO ParentChild algorithm (Grossmann et al. 2007);
vi. the subset of the total significant list for GOParsimony (precedence≤3);
vii. the subset of the total significant list for GOParsimony (precedence≤5);
viii. the subset of the total significant list that intersects with the list produced by the TopGo w1 algorithm (Alexa et al. 2006). This algorithm does not apply a correction for multiple hypotheses and produces a much longer list of GO terms.
ix. the subset of the total significant list that intersects with the list produced by the ShinyGO webserver (Ge et al. 2020). This algorithm applies its own liberal correction for removing terms for >95% similar ‘pathways’. It produces a much longer list of GO terms.

Sets (vi) and (vii) are included which involve higher GOParsimony precedence level thresholds, to see whether they are similar to sets from any of the other diverse methods, but conceptually their usage is very different; they comprise greater GO term detail that is typically clustered near terms from the MP precedence=1 set. Also, for sets (viii) and (ix) the values of the total % of the total significant list that they cover, or the mean –log[P-value], do not include all the other less significant terms that appear in their outputs. Their whole lists of terms are however used to calculate the method clustering.

In general, the most parsimonious precedence=1 GOParsimony set has greatest mean significance (higher mean –log[P-value]). For the three yeast sets, it contains 12-24% of the total significant list, whereas some other methods consistently cover less of it (GOSlims and TopGO ParentChild, 5-18%), and others more (default Revigo, 51-82%). For the two human data sets, the most parsimonious precedence=1 GOParsimony sets cover only 2% of the total significant list, whereas all the other methods cover greater percentages, up to 55-56% for default Revigo). This lower human GOParsimony coverage may simply be a consequence of more detailed functional annotation that is distributed across human proteins/genes.

## Discussion

This tool applies a novel algorithm for reducing GO term redundancy. The basic idea is to apply Occam’s razor (deriving the simplest explanation for phenomena). GOParsimony spits out the most parsimonious (i.e., most simply explanatory) set, assigning it precedence rank = 1. Other precedence ranks are assigned using cascades of precedence incrementation that rapidly converge. These higher precedence ranks can be investigated for insights into functional modules, particularly through perusal of the 2D heatmap plot that is output. In this heatmap, as can be seen in the five examples, GO terms from the three different parts of the digraph (cellular component, molecular function, biological process) can be clustered together, which is advantageous for further hypothesis generation.

In the examples, it was observed that an inherent property of the GO digraph design is that certain more general terms are prolific in terms of immediate children, *e*.*g*., *protein binding* or *protein-containing complex*. When such terms are assigned precedence =1, such diverse children arrays can easily be displayed by setting the maximum precedence level to capture them (usually =2 or =3).

Because the output figures are potentially oversized for very large data sets, to preserve sanity, users may wish to restrict the output to simply drawing the precedence=1 sets. Alternatively, the user can ‘zoom in’ to particular favoured graph or plot areas, depending on whatever further hypotheses are being generated.

## Conclusions

Every year across the planet, thousands of biological researchers become ontically overwhelmed by the functional implications of their data, as presented to them through the lens of Gene Ontology. GOParsimony applies the principle of parsimony—that is, aspiring to the simplest explanation for phenomena—to derive a single concise non-redundant list of GO terms for a protein/gene data set, while also allowing for the discovery of functional modules that might trigger further hypothesizing. It achieves this by saltating across the GO digraph, releasing cascades of GO term precedence in the process. For five example data sets discussed, GOParsimony consistently produced reduced GO term lists of greater significance and low to intermediate coverage of the space of significant GO terms. GOParsimony can be readily grafted into investigators’ computational pipelines.

