## Supplementary Files 1 to 5 for "*GOParsimony*: parsimonious saltation across the Gene Ontology digraph for efficient functional annotation": Suppl-File-1-Example-1-yeast-AQrich-cluster.pdf

#### Suppl. File 1: Graphical GOParsimony output for Example 1, the yeast AQ-rich intrinsically-disordered cluster

**Figure A: Complete precedence graph (default precedence <=5).** The GO terms are labelled in the upper right corner with their precedence levels and are colour-graded with a blueness scale. The GO-term boxes also contain the term definition, the corrected P-value (Q-value if submitted to the Benjamini-Hochberg procedure), and the term type in the lower right corner (BP for biological process, CC for cellular component, MF for molecular function). They are connected with dotted arrows if there are only intervening non-significant GO terms. GOslim terms are outlined in magenta, and the user-specified highlight list in green. Here, the highlights come from default labelling of non-redundant GO terms with the Revigo website.

**Figure B: Concise precedence graph (default precedence <=5).** It is the same as in Figure A, except that only terms with precedence levels ≤ the level user-specified are included (here, the default = 5 is used).

**Figure C: 2D Heatmap for inferring functional modules. Proteins**/genes are clustered on the y-axis using a Jaccard distance calculated from the common GO terms. GO terms are clustered on the x-axis using a Jaccard distance calculated from the common proteins/genes. GO term names are colour-coded according to type (brown = CC, yellow = BP, green = MF). Cells are colour-coded for precedence as in the precedence graphs, and also labelled with the precedence level.

**Figure D: Comparison graph bar-chart indicating GO terms added as children or parents for higher precedence level thresholds.** Precedence level thresholds up to the precedence level specified in the GOParsimony run are shown (here, the default level ≤5).

**Figure E: Comparison graph Venn diagram.** The overlaps in the Precedence=1, default Revigo and GOslims term sets are shown. The table lists the percentage coverage of significant terms and the average –log(P-value) (i.e., corrected P-value).

**Figure F: Coloured GO term table.** It is colour-coded to correspond to the precedence graphs.

### PRECEDENCE GRAPHS

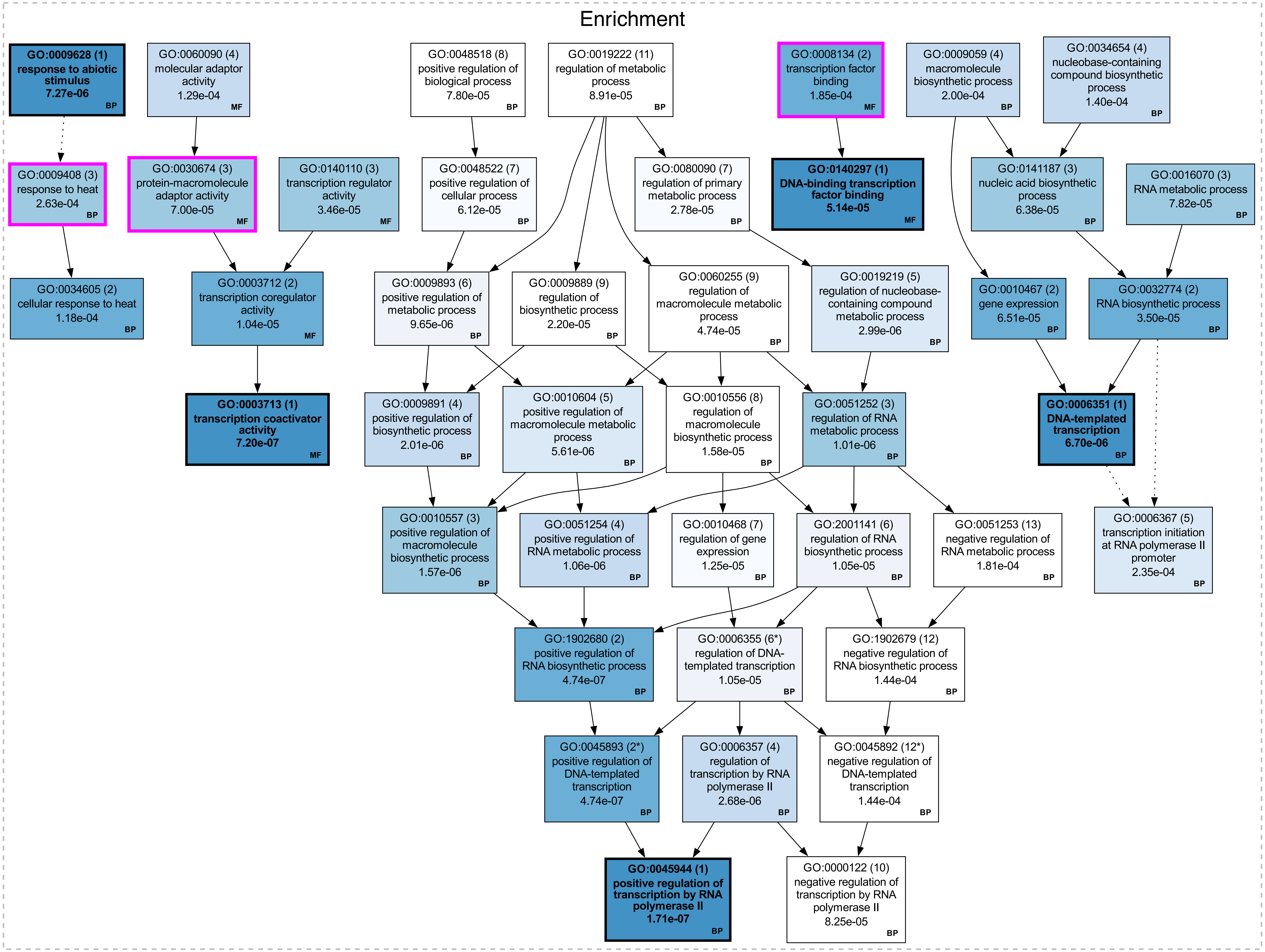

CONCISE PRECEDENCE GRAPHS (Prec <= 5)

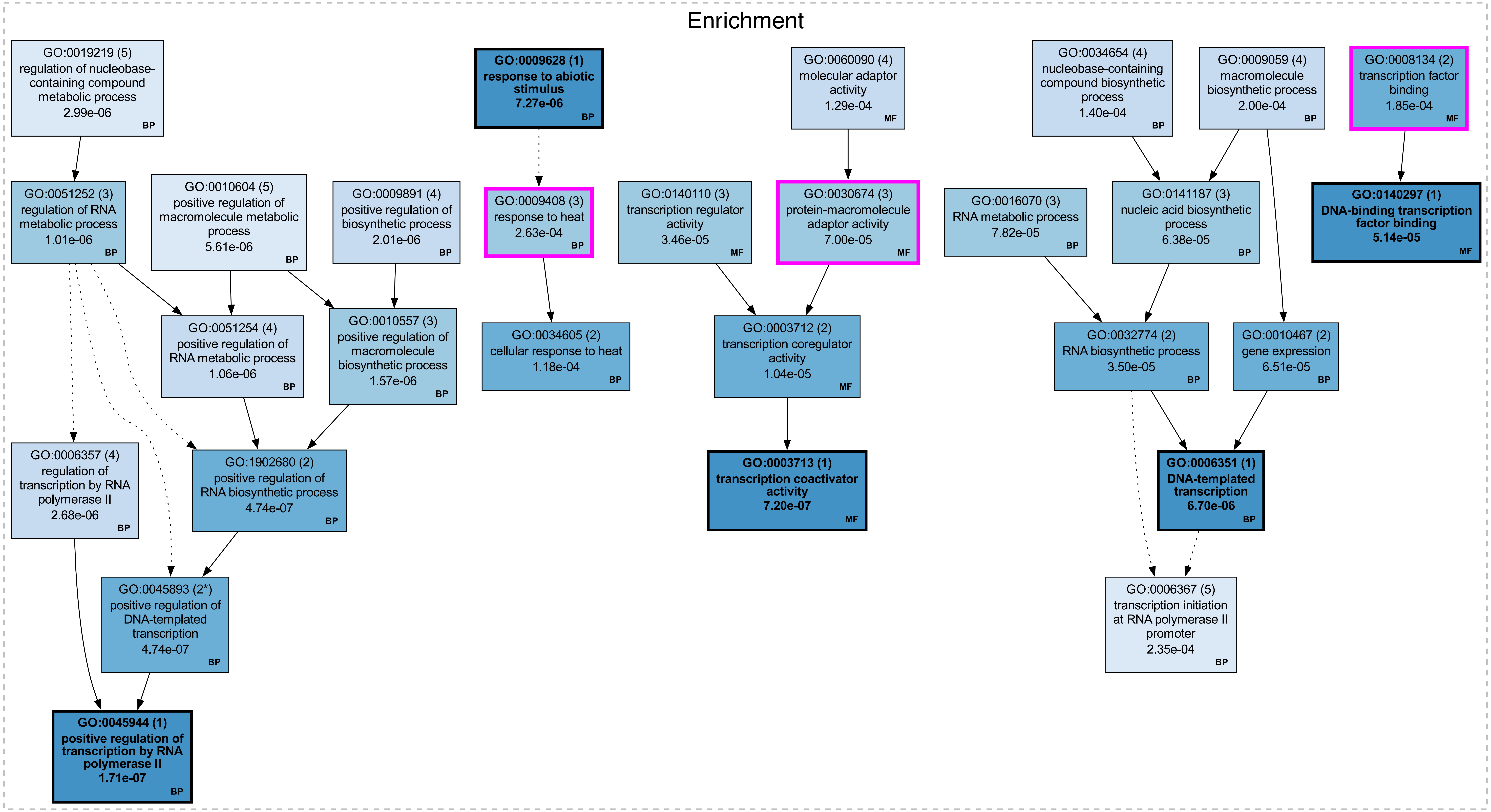

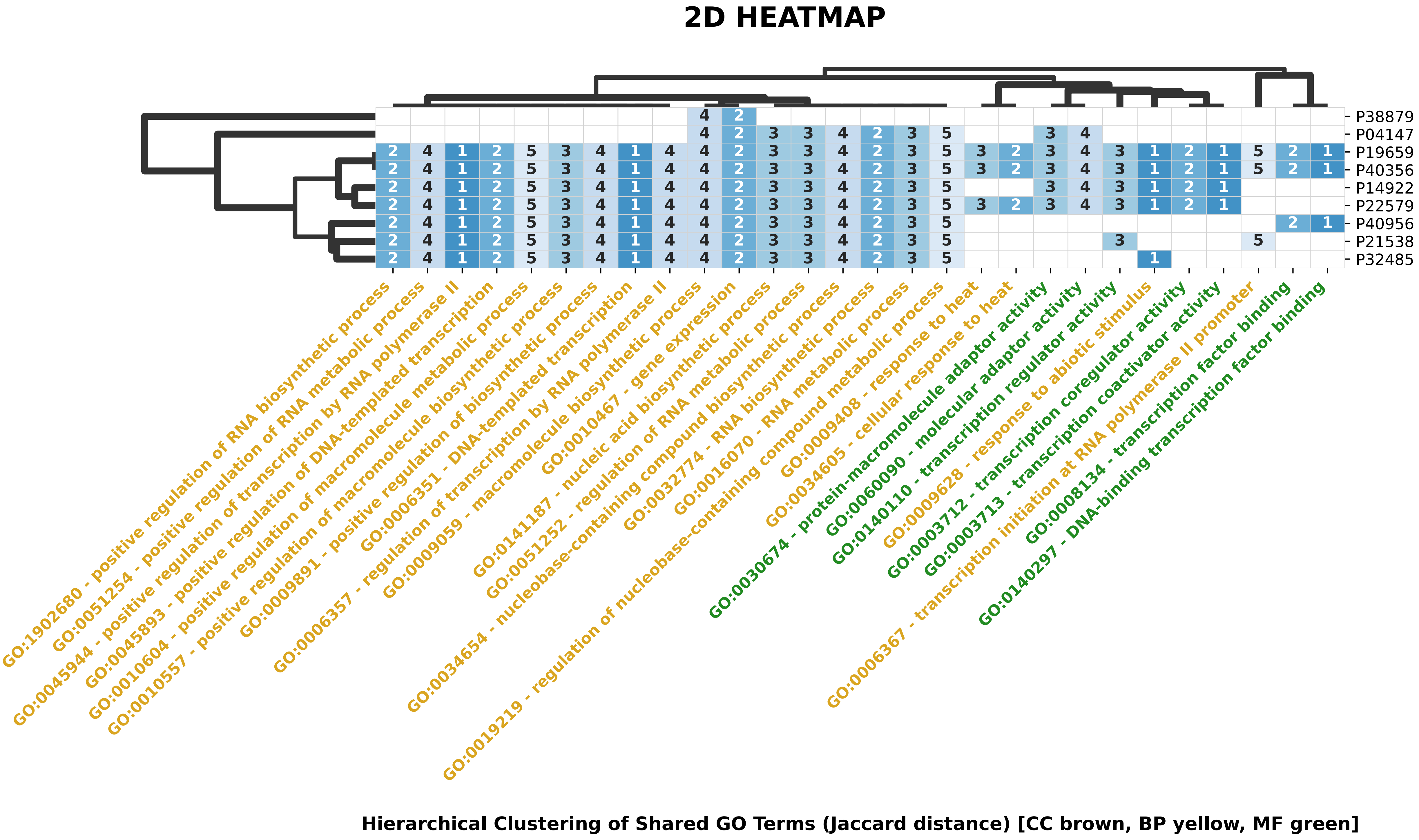
2D HEATMAP
P38879
P04147
P19659
P40356
P14922
P22579
P40956
P21538
P32485
GO:1902680 - positive regulation of RNA biosynthetic process
GO:0051254 - positive regulation of RNA metabolic process
GO:0045944 - positive regulation of transcription by RNA polymerase II
GO:0045893 - positive regulation of DNA-templated transcription
GO:0010604 - positive regulation of macromolecule metabolic process
GO:0010557 - positive regulation of macromolecule biosynthetic process
GO:0009891 - positive regulation of biosynthetic process
GO:0006351 - DNA-templated transcription
GO:0006357 - regulation of transcription by RNA polymerase II
GO:0009059 - macromolecule biosynthetic process
GO:0010467 - gene expression
GO:0141187 - nucleic acid biosynthetic process
GO:0051252 - regulation of RNA metabolic process
GO:0034654 - nucleobase-containing compound biosynthetic process
GO:0032774 - RNA biosynthetic process
GO:0016070 - RNA metabolic process
GO:0019219 - regulation of nucleobase-containing compound metabolic process
GO:0009408 - response to heat
GO:0034605 - cellular response to heat
GO:0030674 - protein-macromolecule adaptor activity
GO:0060090 - molecular adaptor activity
GO:0140110 - transcription regulator activity
GO:0009628 - response to abiotic stimulus
GO:0003712 - transcription coregulator activity
GO:0003713 - transcription coactivator activity
GO:0006367 - transcription initiation at RNA polymerase II promoter
GO:0008134 - transcription factor binding
GO:0140297 - DNA-binding transcription factor binding
Hierarchical Clustering of Shared GO Terms (Jaccard distance) [CC brown, BP yellow, MF green]

### COMPARISON GRAPHS

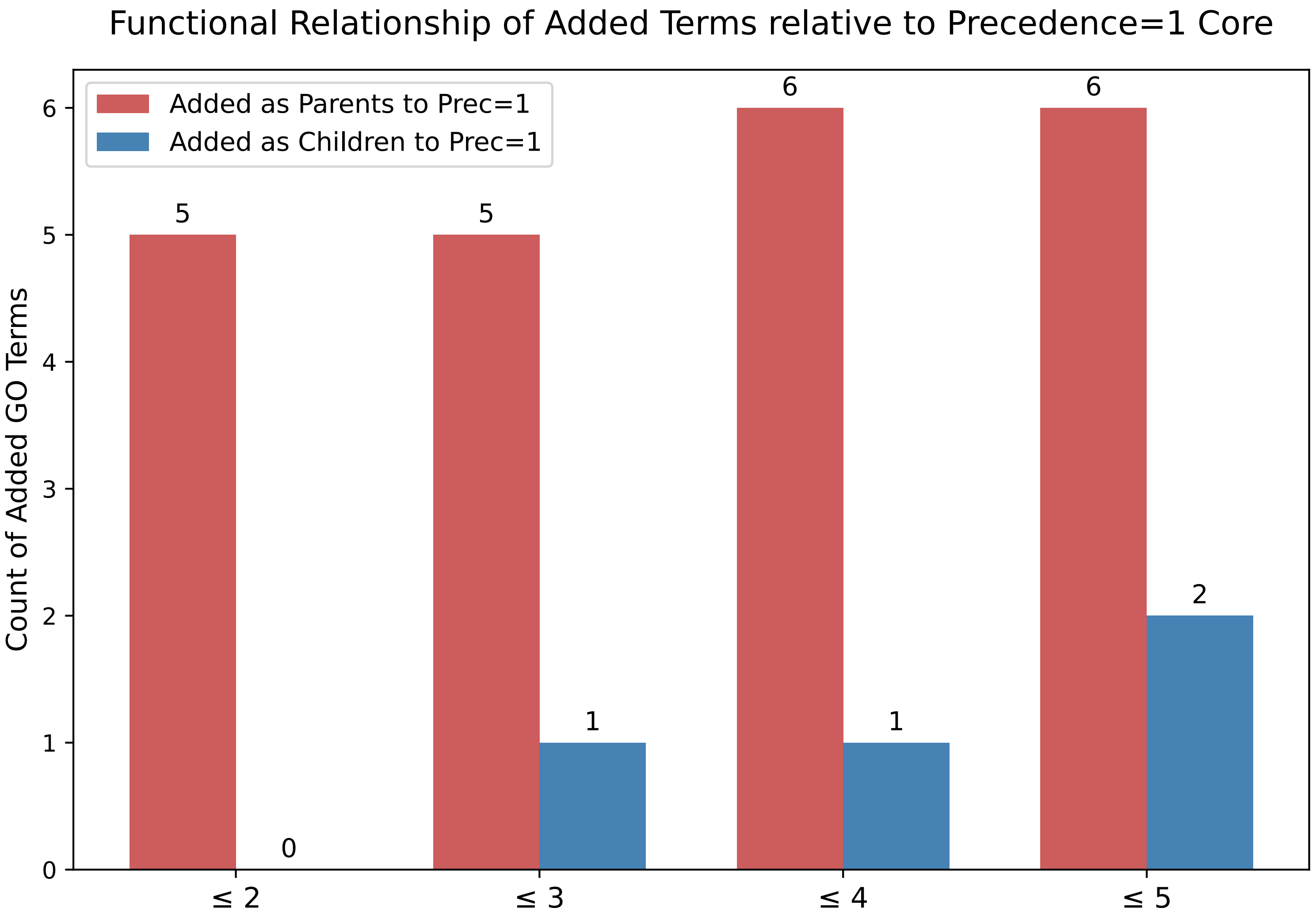

Overlap of Significant GO Terms

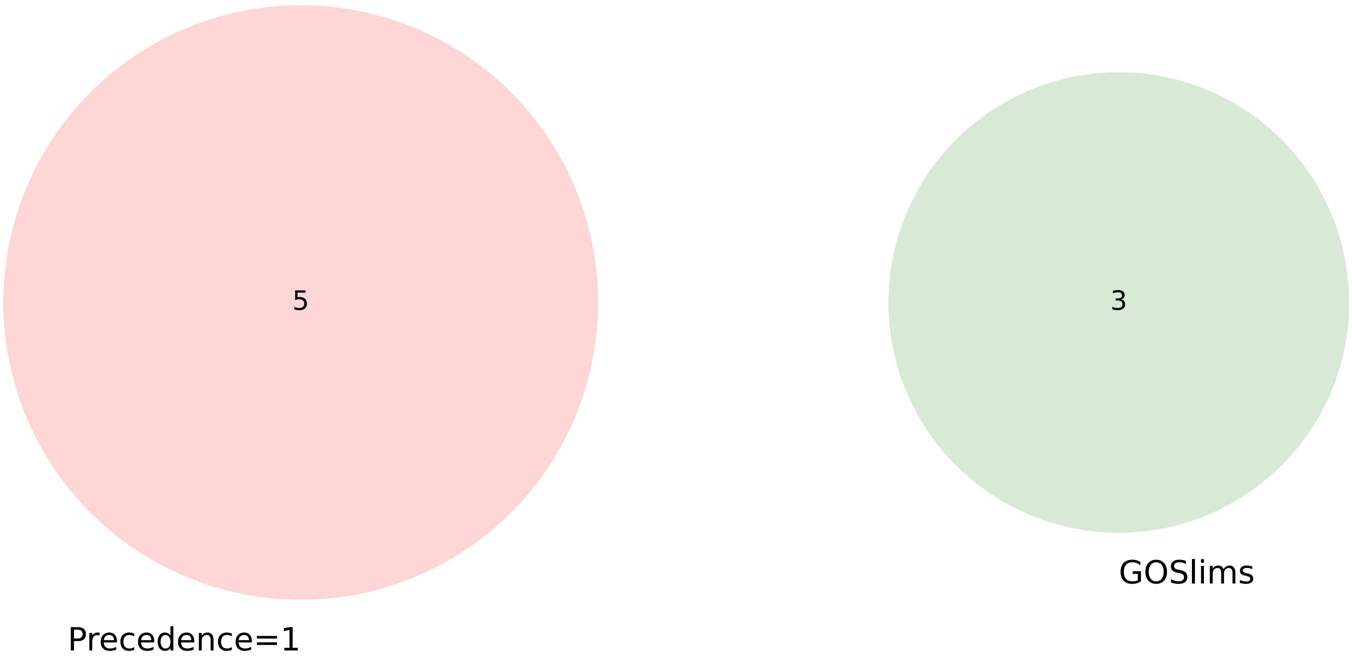


| Set Name | % Coverage (of the significant terms) | Average -log10(P-value) |
|---|---|---|
| **Precedence=1** | 11.6% | 5.50 |
| **GOSlims** | 7.0% | 3.82 |

### GOTERM TABLE (Prec <= 5)

| Namespace | P-Value | Enr/Dep | Prec | Reg_Prec | Reg_ClstID | Slim | GOTerm | Description |
|---|---|---|---|---|---|---|---|---|
| BP | 1.71e-07 | Enr | 1 | 1 | 0 | No | GO:0045944 | positive regulation of transcription by RNA polymerase II |
| BP | 4.74e-07 | Enr | 2* | 1 | 9 | No | GO:0045893 | positive regulation of DNA-templated transcription |
| BP | 4.74e-07 | Enr | 2 | 1 | 10 | No | GO:1902680 | positive regulation of RNA biosynthetic process |
| MF | 7.20e-07 | Enr | 1 | 1 | 0 | No | GO:0003713 | transcription coactivator activity |
| BP | 1.01e-06 | Enr | 3 | 1 | 11 | No | GO:0051252 | regulation of RNA metabolic process |
| BP | 1.06e-06 | Enr | 4 | 1 | 11 | No | GO:0051254 | positive regulation of RNA metabolic process |
| BP | 1.57e-06 | Enr | 3 | 1 | 12 | No | GO:0010557 | positive regulation of macromolecule biosynthetic process |
| BP | 2.01e-06 | Enr | 4 | 1 | 0 | No | GO:0009891 | positive regulation of biosynthetic process |
| BP | 2.68e-06 | Enr | 4 | 1 | 0 | No | GO:0006357 | regulation of transcription by RNA polymerase II |
| BP | 2.99e-06 | Enr | 5 | 1 | 0 | No | GO:0019219 | regulation of nucleobase-containing compound metabolic process |
| BP | 5.61e-06 | Enr | 5 | 1 | 0 | No | GO:0010604 | positive regulation of macromolecule metabolic process |
| BP | 6.70e-06 | Enr | 1 | 2 | 9 | No | GO:0006351 | DNA-templated transcription |
| BP | 7.27e-06 | Enr | 1 | 1 | 0 | No | GO:0009628 | response to abiotic stimulus |
| MF | 1.04e-05 | Enr | 2 | 1 | 0 | No | GO:0003712 | transcription coregulator activity |
| MF | 3.46e-05 | Enr | 3 | 1 | 0 | No | GO:0140110 | transcription regulator activity |
| BP | 3.50e-05 | Enr | 2 | 2 | 10 | No | GO:0032774 | RNA biosynthetic process |
| MF | 5.14e-05 | Enr | 1 | 1 | 0 | No | GO:0140297 | DNA-binding transcription factor binding |
| BP | 6.38e-05 | Enr | 3 | 1 | 0 | No | GO:0141187 | nucleic acid biosynthetic process |
| BP | 6.51e-05 | Enr | 2 | 2 | 19 | No | GO:0010467 | gene expression |
| MF | 7.00e-05 | Enr | 3 | 1 | 0 | Yes | GO:0030674 | protein-macromolecule adaptor activity |
| BP | 7.82e-05 | Enr | 3 | 2 | 11 | No | GO:0016070 | RNA metabolic process |
| BP | 1.18e-04 | Enr | 2 | 1 | 0 | No | GO:0034605 | cellular response to heat |
| MF | 1.29e-04 | Enr | 4 | 1 | 0 | No | GO:0060090 | molecular adaptor activity |
| BP | 1.40e-04 | Enr | 4 | 1 | 0 | No | GO:0034654 | nucleobase-containing compound biosynthetic process |
| MF | 1.85e-04 | Enr | 2 | 1 | 0 | Yes | GO:0008134 | transcription factor binding |
| BP | 2.00e-04 | Enr | 4 | 2 | 12 | No | GO:0009059 | macromolecule biosynthetic process |
| BP | 2.35e-04 | Enr | 5 | 1 | 0 | No | GO:0006367 | transcription initiation at RNA polymerase II promoter |
| BP | 2.63e-04 | Enr | 3 | 1 | 0 | Yes | GO:0009408 | response to heat |
