## Supplementary Files 1 to 5 for "*GOParsimony*: parsimonious saltation across the Gene Ontology digraph for efficient functional annotation": Suppl-File-2-Example-2-yeast-EKrich-low-blockiness.pdf

**Figure F: Coloured GO term table.** It is colour-coded to correspond to the precedence graphs.

### PRECEDENCE GRAPHS

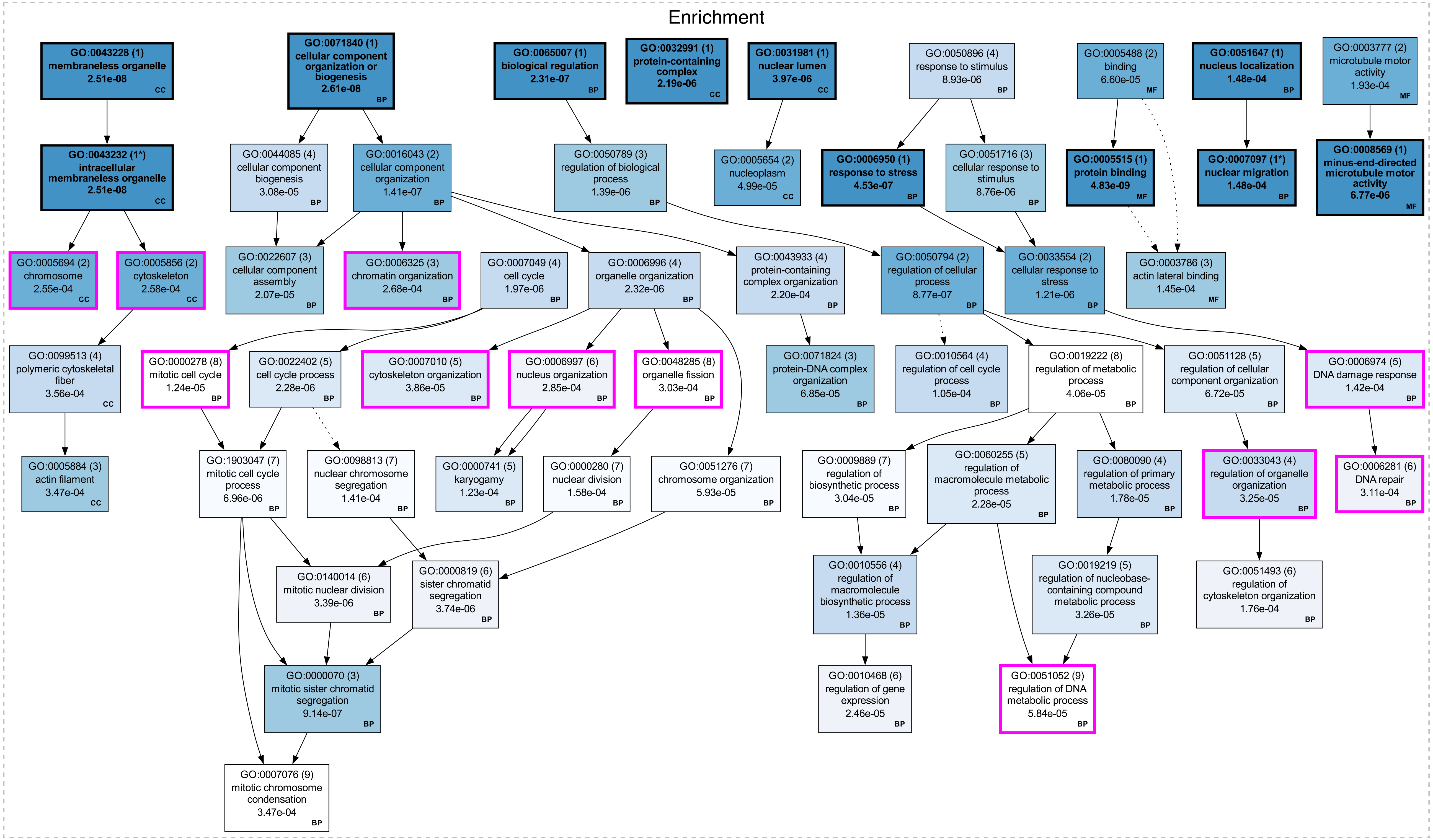

### CONCISE PRECEDENCE GRAPHS (Prec <= 5)

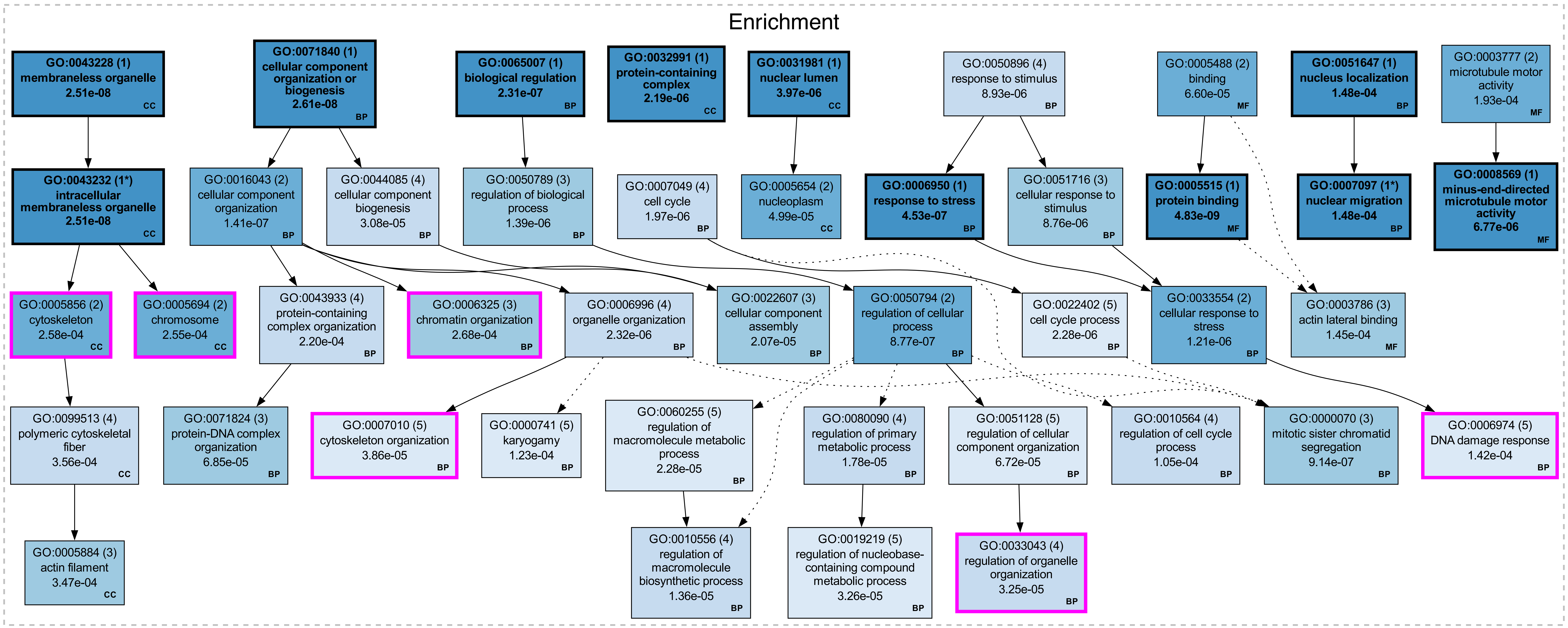

### 2D HEATMAP

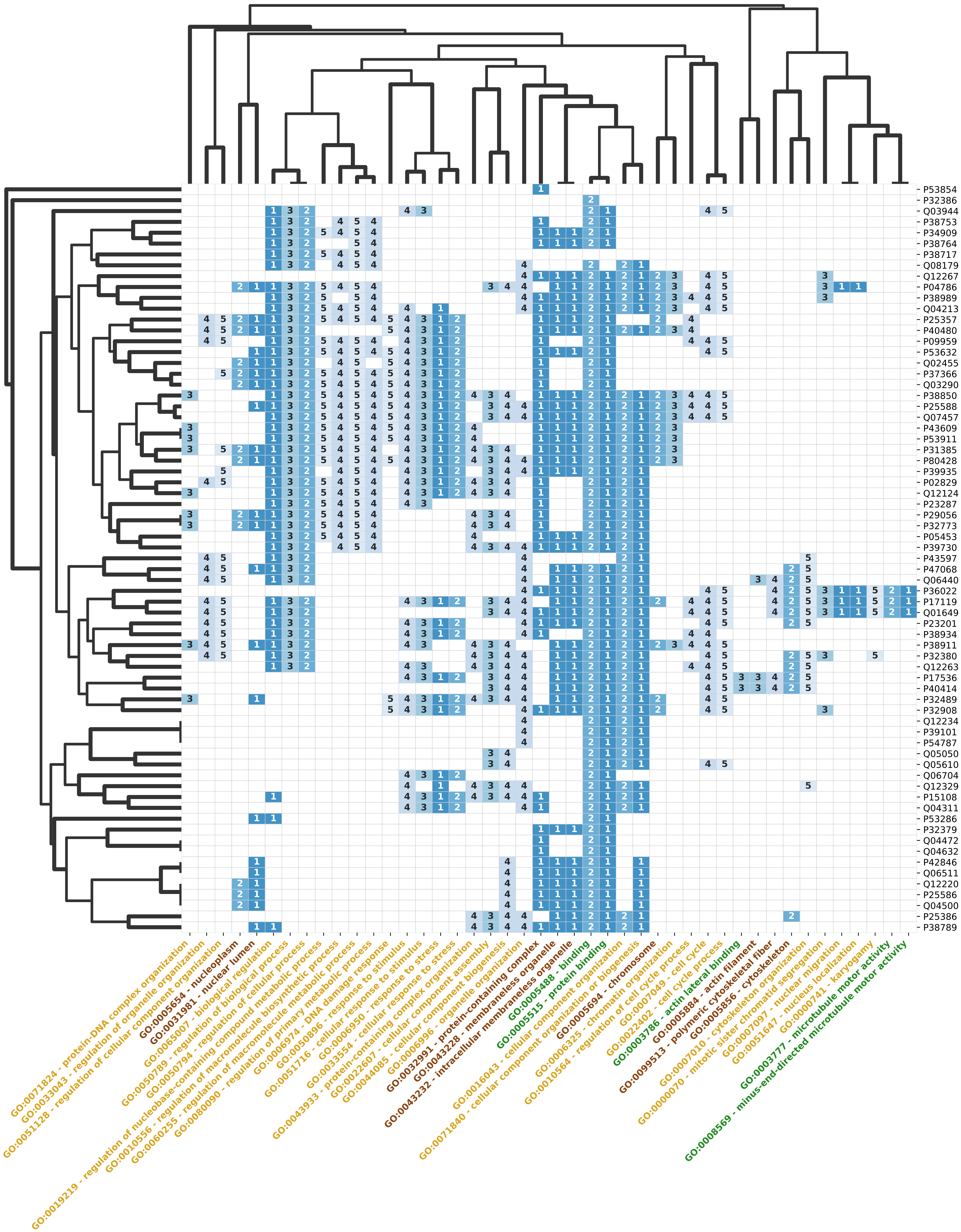


**Hierarchical Clustering of Shared GO Terms (Jaccard distance) [CC brown, BP yellow, MF green]**

### COMPARISON GRAPHS

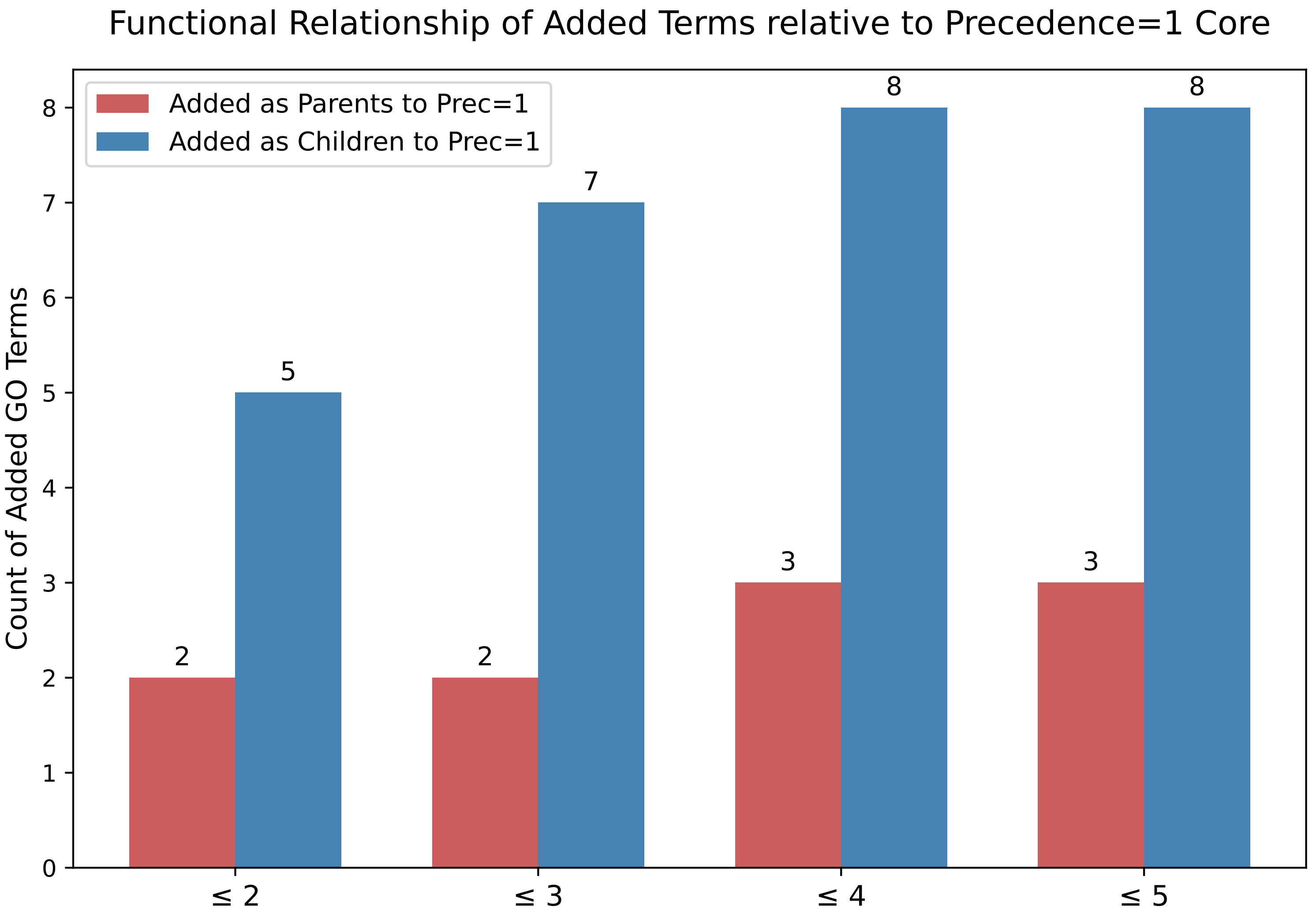

### Overlap of Significant GO Terms

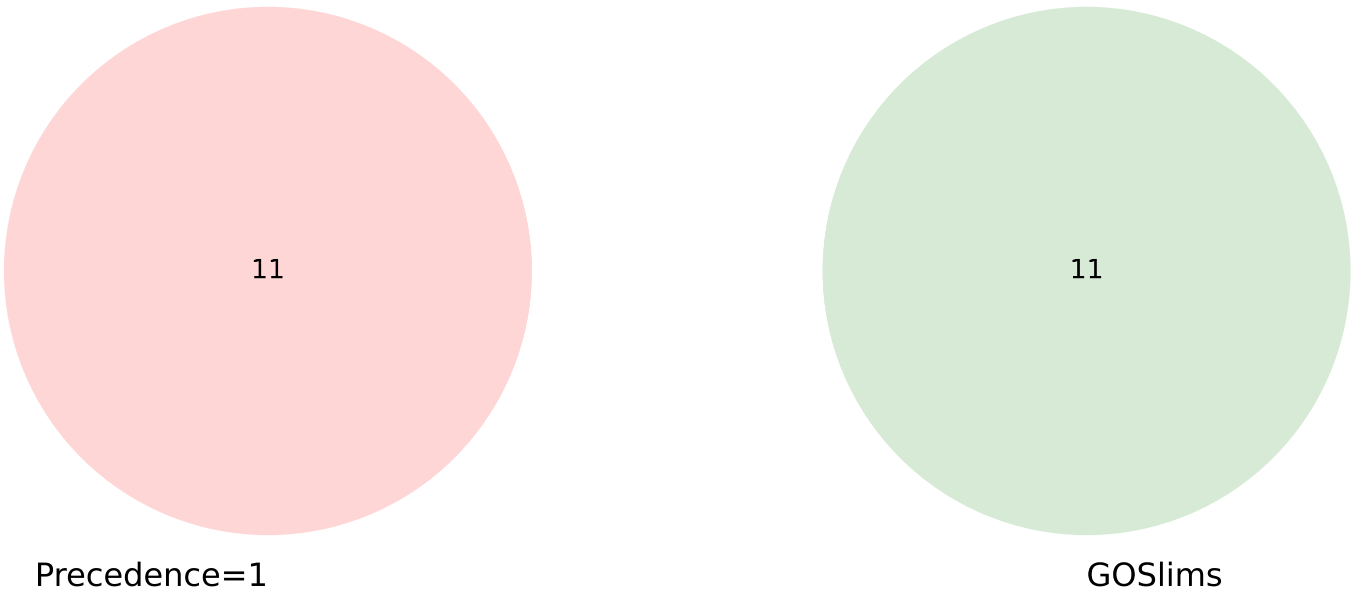


| Set Name | % Coverage (of the significant terms) | Average -log10(P-value) |
|---|---|---|
| **Precedence=1** | 18.3% | 6.18 |
| **GOSlims** | 18.3% | 3.93 |

### GOTERM TABLE (Prec <= 5)

| Namespace | P-Value | Enr/Dep | Prec | Reg_Prec | Reg_ClstID | Slim | GOTerm | Description |
|---|---|---|---|---|---|---|---|---|
| MF | 4.83e-09 | Enr | 1 | 1 | 0 | No | GO:0005515 | protein binding |
| CC | 2.51e-08 | Enr | 1* | 1 | 0 | No | GO:0043232 | intracellular membraneless organelle |
| CC | 2.51e-08 | Enr | 1 | 1 | 0 | No | GO:0043228 | membraneless organelle |
| BP | 2.61e-08 | Enr | 1 | 1 | 0 | No | GO:0071840 | cellular component organization or biogenesis |
| BP | 1.41e-07 | Enr | 2 | 1 | 13 | No | GO:0016043 | cellular component organization |
| BP | 2.31e-07 | Enr | 1 | 1 | 0 | No | GO:0065007 | biological regulation |
| BP | 4.53e-07 | Enr | 1 | 1 | 0 | No | GO:0006950 | response to stress |
| BP | 8.77e-07 | Enr | 2 | 1 | 0 | No | GO:0050794 | regulation of cellular process |
| BP | 9.14e-07 | Enr | 3 | 1 | 0 | No | GO:0000070 | mitotic sister chromatid segregation |
| BP | 1.21e-06 | Enr | 2 | 1 | 0 | No | GO:0033554 | cellular response to stress |
| BP | 1.39e-06 | Enr | 3 | 1 | 0 | No | GO:0050789 | regulation of biological process |
| BP | 1.97e-06 | Enr | 4 | 1 | 0 | No | GO:0007049 | cell cycle |
| CC | 2.19e-06 | Enr | 1 | 1 | 0 | No | GO:0032991 | protein-containing complex |
| BP | 2.28e-06 | Enr | 5 | 1 | 21 | No | GO:0022402 | cell cycle process |
| BP | 2.32e-06 | Enr | 4 | 1 | 22 | No | GO:0006996 | organelle organization |
| CC | 3.97e-06 | Enr | 1 | 1 | 0 | No | GO:0031981 | nuclear lumen |
| MF | 6.77e-06 | Enr | 1 | 1 | 0 | No | GO:0008569 | minus-end-directed microtubule motor activity |
| BP | 8.76e-06 | Enr | 3 | 1 | 0 | No | GO:0051716 | cellular response to stimulus |
| BP | 8.93e-06 | Enr | 4 | 1 | 0 | No | GO:0050896 | response to stimulus |
| BP | 1.36e-05 | Enr | 4 | 1 | 0 | No | GO:0010556 | regulation of macromolecule biosynthetic process |
| BP | 1.78e-05 | Enr | 4 | 1 | 0 | No | GO:0080090 | regulation of primary metabolic process |
| BP | 2.07e-05 | Enr | 3 | 1 | 0 | No | GO:0022607 | cellular component assembly |
| BP | 2.28e-05 | Enr | 5 | 1 | 0 | No | GO:0060255 | regulation of macromolecule metabolic process |
| BP | 3.08e-05 | Enr | 4 | 1 | 0 | No | GO:0044085 | cellular component biogenesis |
| BP | 3.25e-05 | Enr | 4 | 2 | 22 | Yes | GO:0033043 | regulation of organelle organization |
| BP | 3.26e-05 | Enr | 5 | 1 | 0 | No | GO:0019219 | regulation of nucleobase-containing compound metabolic process |
| BP | 3.86e-05 | Enr | 5 | 1 | 37 | Yes | GO:0007010 | cytoskeleton organization |
| CC | 4.99e-05 | Enr | 2 | 1 | 0 | No | GO:0005654 | nucleoplasm |
| MF | 6.60e-05 | Enr | 2 | 1 | 0 | No | GO:0005488 | binding |
| BP | 6.72e-05 | Enr | 5 | 2 | 13 | No | GO:0051128 | regulation of cellular component organization |
| BP | 6.85e-05 | Enr | 3 | 1 | 0 | No | GO:0071824 | protein-DNA complex organization |
| BP | 1.05e-04 | Enr | 4 | 2 | 21 | No | GO:0010564 | regulation of cell cycle process |
| BP | 1.23e-04 | Enr | 5 | 1 | 0 | No | GO:0000741 | karyogamy |
| BP | 1.42e-04 | Enr | 5 | 1 | 0 | Yes | GO:0006974 | DNA damage response |
| MF | 1.45e-04 | Enr | 3 | 1 | 0 | No | GO:0003786 | actin lateral binding |
| BP | 1.48e-04 | Enr | 1* | 1 | 0 | No | GO:0007097 | nuclear migration |
| BP | 1.48e-04 | Enr | 1 | 1 | 0 | No | GO:0051647 | nucleus localization |
| MF | 1.93e-04 | Enr | 2 | 1 | 0 | No | GO:0003777 | microtubule motor activity |
| BP | 2.20e-04 | Enr | 4 | 1 | 0 | No | GO:0043933 | protein-containing complex organization |
| CC | 2.55e-04 | Enr | 2 | 1 | 0 | Yes | GO:0005694 | chromosome |
| CC | 2.58e-04 | Enr | 2 | 1 | 0 | Yes | GO:0005856 | cytoskeleton |
| BP | 2.68e-04 | Enr | 3 | 1 | 0 | Yes | GO:0006325 | chromatin organization |
| CC | 3.47e-04 | Enr | 3 | 1 | 0 | No | GO:0005884 | actin filament |
| CC | 3.56e-04 | Enr | 4 | 1 | 0 | No | GO:0099513 | polymeric cytoskeletal fiber |
