## Supplementary Files 1 to 5 for "*GOParsimony*: parsimonious saltation across the Gene Ontology digraph for efficient functional annotation": Suppl-File-3-Example-3-yeast-EKrich-high-blockiness.pdf

**Figure F: Coloured GO term table.** It is colour-coded to correspond to the precedence graphs.

#### PRECEDENCE GRAPHS

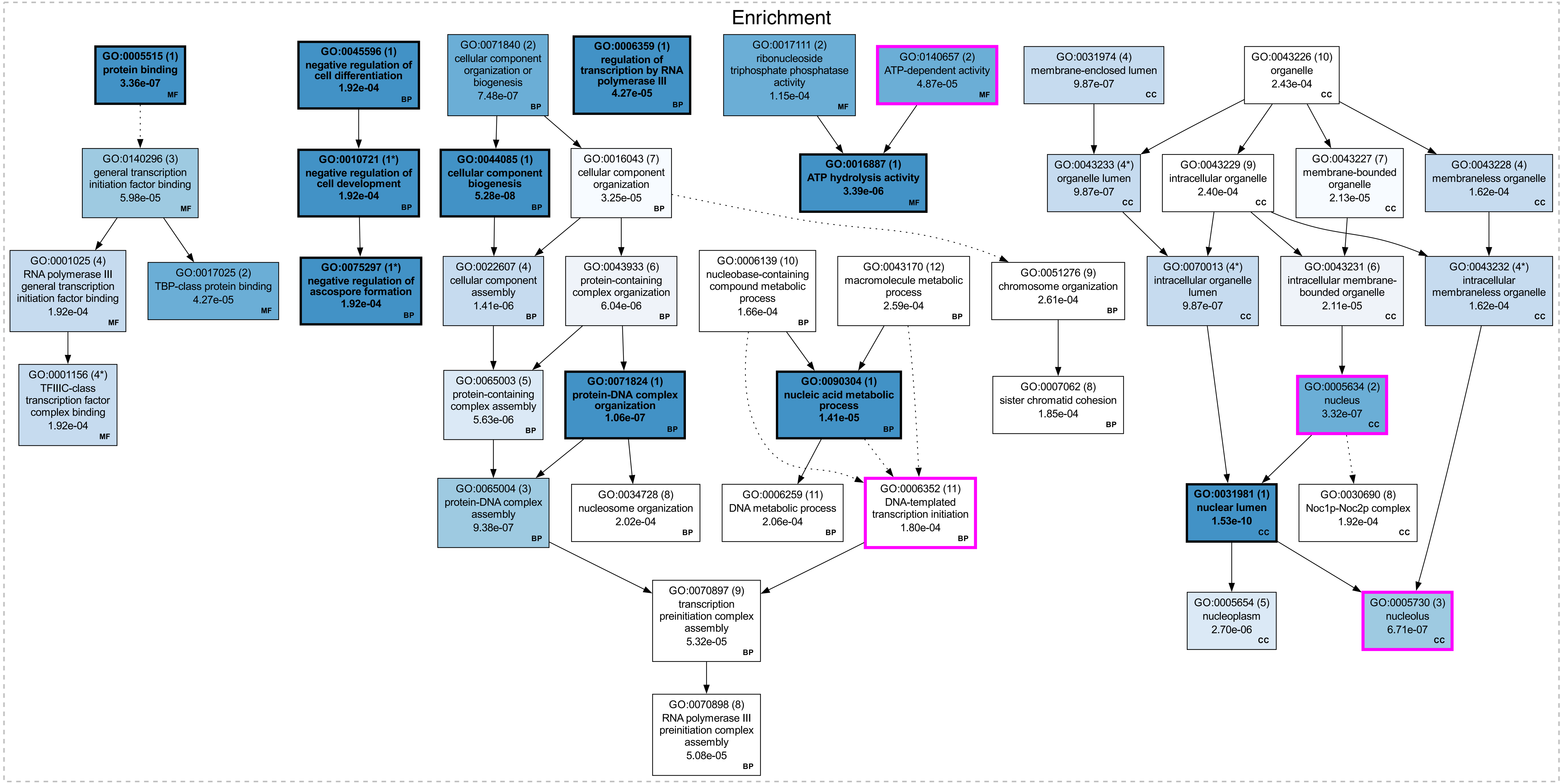

CONCISE PRECEDENCE GRAPHS (Prec <= 5)

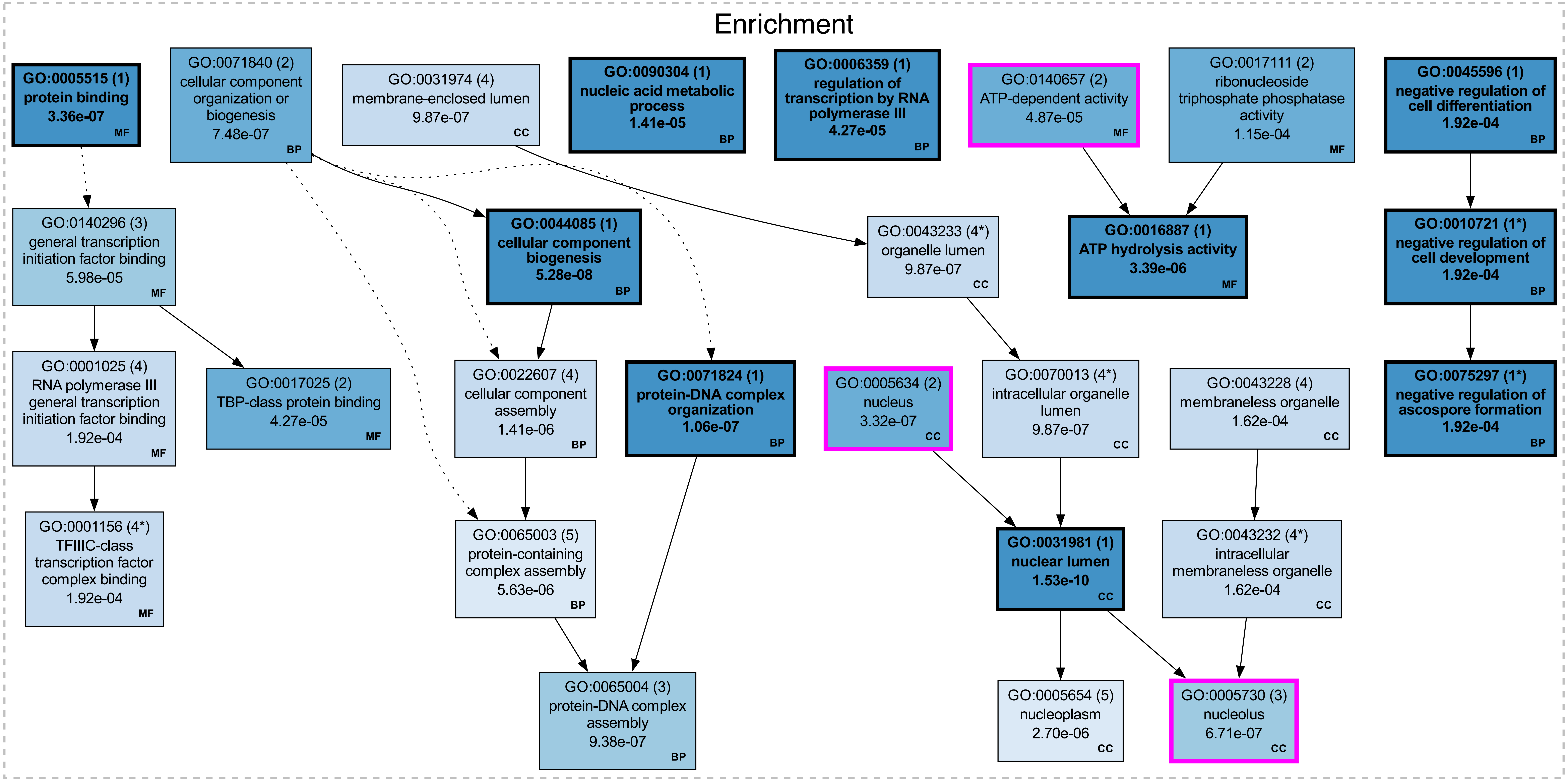

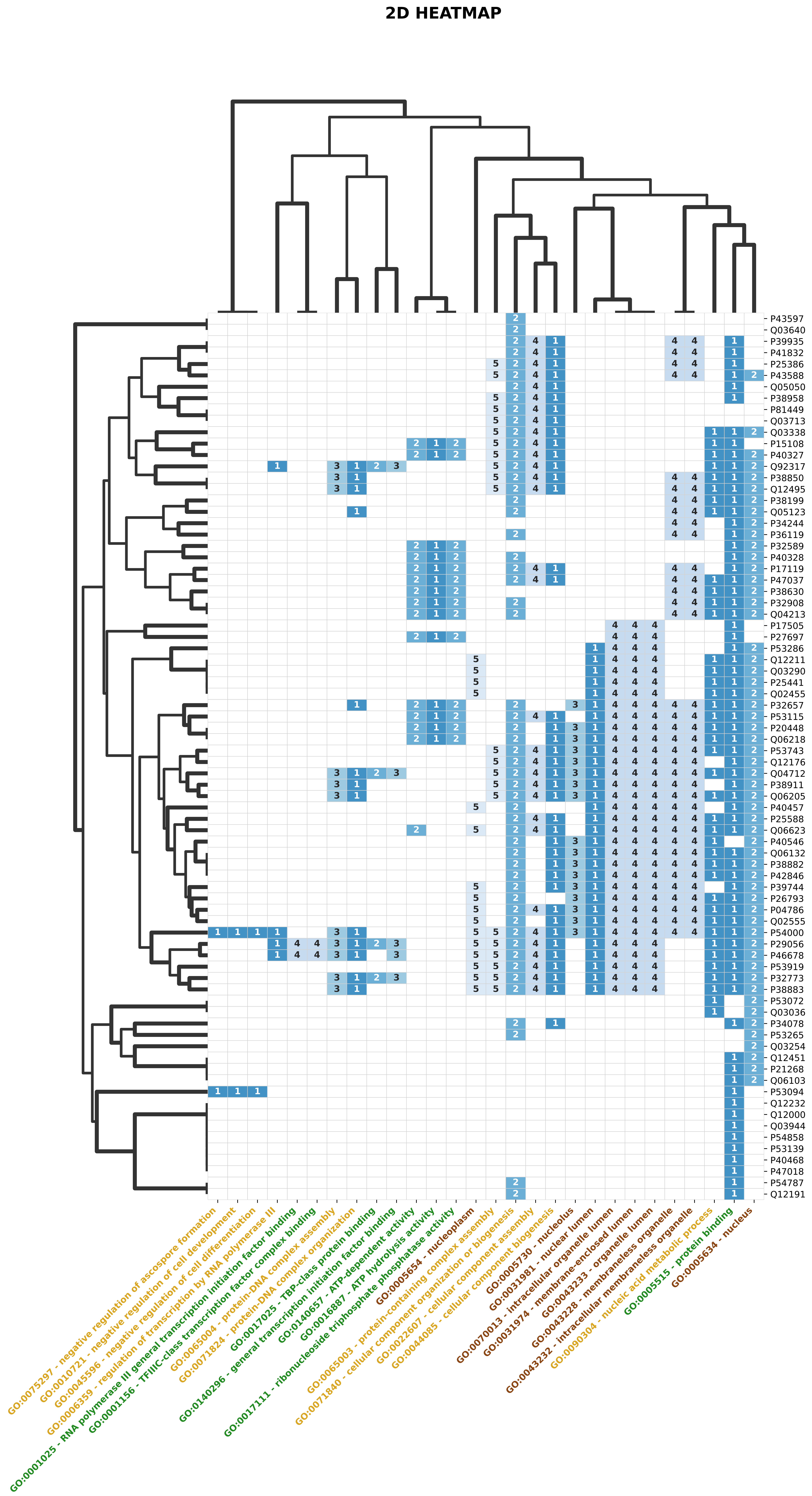


Hierarchical Clustering of Shared GO Terms (Jaccard distance) [CC brown, BP yellow, MF green]

### COMPARISON GRAPHS

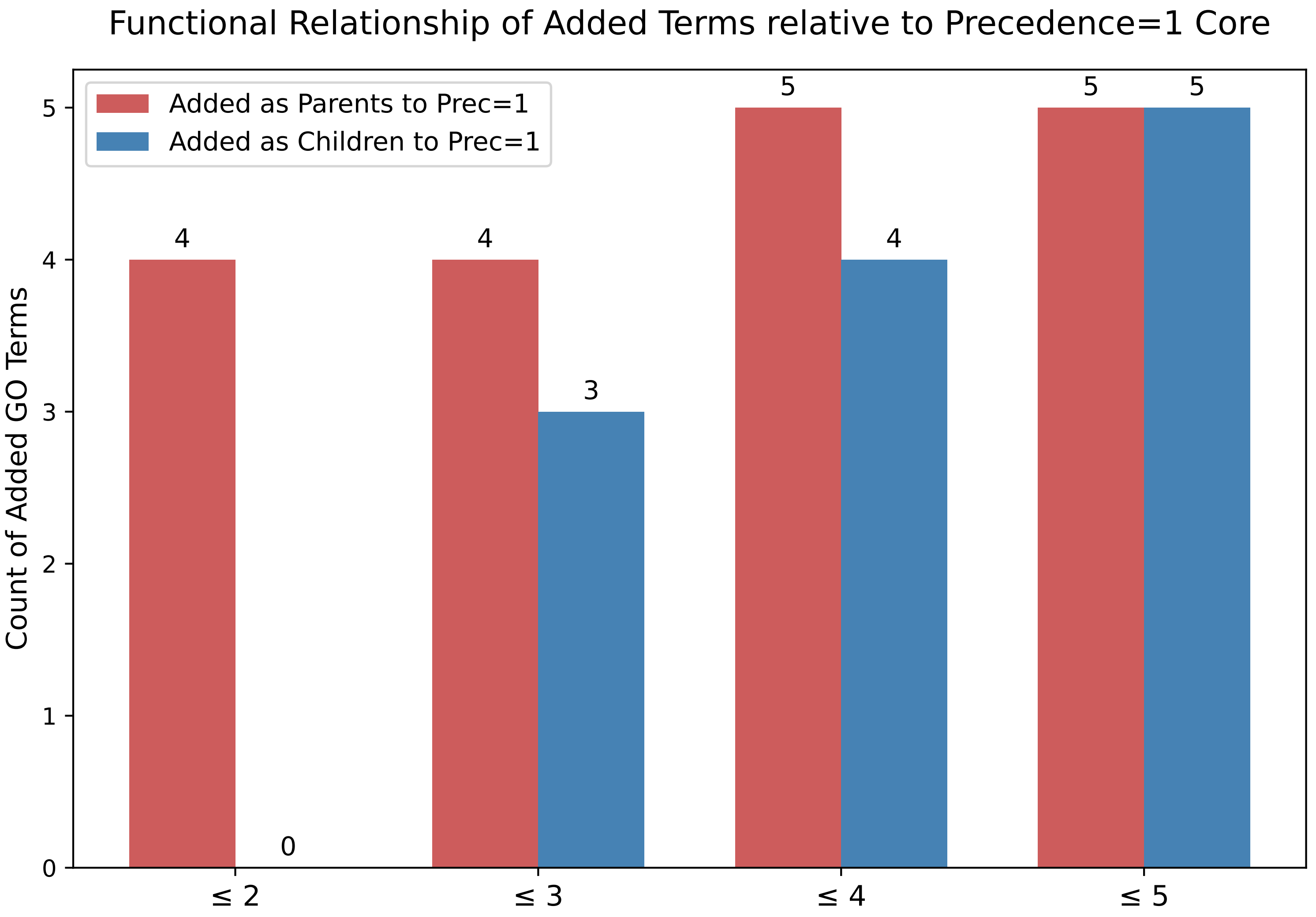

Overlap of Significant GO Terms

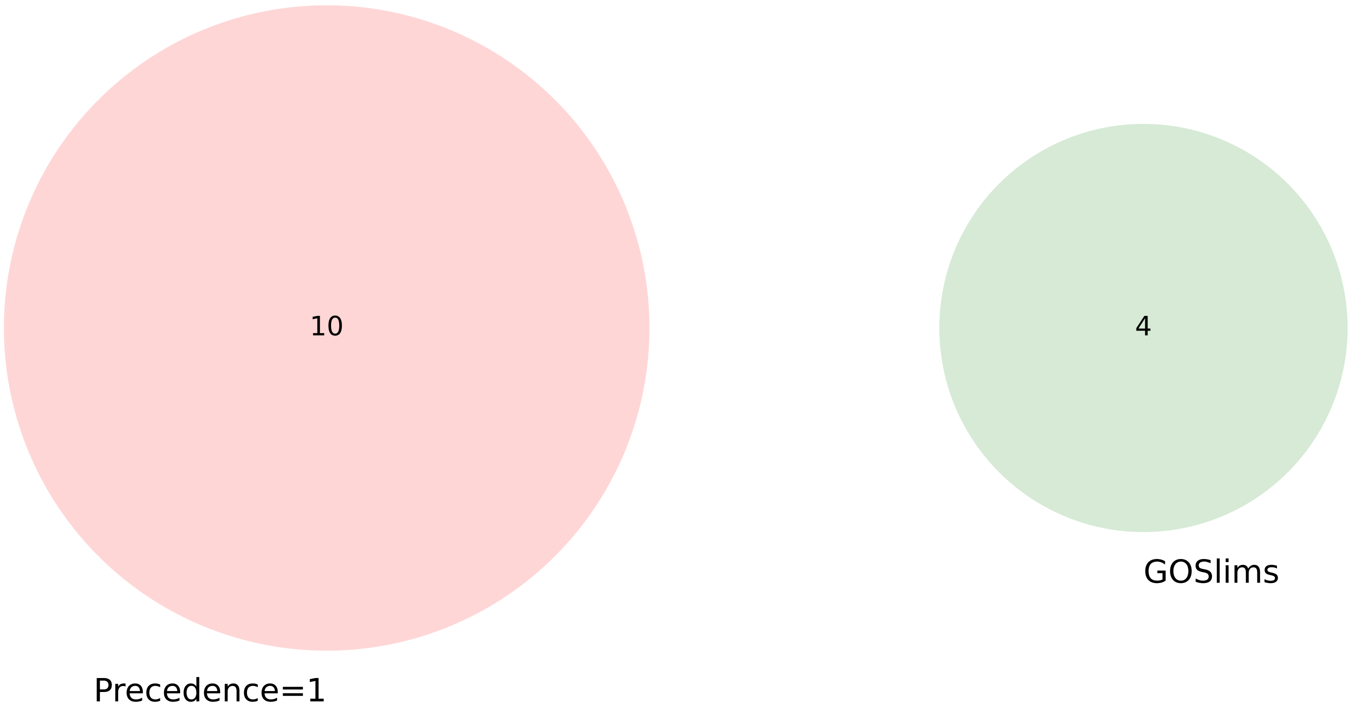


| Set Name | % Coverage (of the significant terms) | Average -log10(P-value) |
|---|---|---|
| **Precedence=1** | 22.7% | 5.64 |
| **GOSlims** | 9.1% | 5.18 |

### GOTERM TABLE (Prec <= 5)

| Namespace | P-Value | Enr/Dep | Prec | Reg_Prec | Reg_ClstID | Slim | GOTerm | Description |
|---|---|---|---|---|---|---|---|---|
| CC | 1.53e-10 | Enr | 1 | 1 | 0 | No | GO:0031981 | nuclear lumen |
| BP | 5.28e-08 | Enr | 1 | 1 | 0 | No | GO:0044085 | cellular component biogenesis |
| BP | 1.06e-07 | Enr | 1 | 1 | 0 | No | GO:0071824 | protein-DNA complex organization |
| CC | 3.32e-07 | Enr | 2 | 1 | 0 | Yes | GO:0005634 | nucleus |
| MF | 3.36e-07 | Enr | 1 | 1 | 0 | No | GO:0005515 | protein binding |
| CC | 6.71e-07 | Enr | 3 | 1 | 0 | Yes | GO:0005730 | nucleolus |
| BP | 7.48e-07 | Enr | 2 | 1 | 0 | No | GO:0071840 | cellular component organization or biogenesis |
| BP | 9.38e-07 | Enr | 3 | 1 | 0 | No | GO:0065004 | protein-DNA complex assembly |
| CC | 9.87e-07 | Enr | 4* | 1 | 0 | No | GO:0070013 | intracellular organelle lumen |
| CC | 9.87e-07 | Enr | 4 | 1 | 0 | No | GO:0031974 | membrane-enclosed lumen |
| CC | 9.87e-07 | Enr | 4* | 1 | 0 | No | GO:0043233 | organelle lumen |
| BP | 1.41e-06 | Enr | 4 | 1 | 0 | No | GO:0022607 | cellular component assembly |
| CC | 2.70e-06 | Enr | 5 | 1 | 0 | No | GO:0005654 | nucleoplasm |
| MF | 3.39e-06 | Enr | 1 | 1 | 0 | No | GO:0016887 | ATP hydrolysis activity |
| BP | 5.63e-06 | Enr | 5 | 1 | 0 | No | GO:0065003 | protein-containing complex assembly |
| BP | 1.41e-05 | Enr | 1 | 1 | 0 | No | GO:0090304 | nucleic acid metabolic process |
| MF | 4.27e-05 | Enr | 2 | 1 | 0 | No | GO:0017025 | TBP-class protein binding |
| BP | 4.27e-05 | Enr | 1 | 1 | 0 | No | GO:0006359 | regulation of transcription by RNA polymerase III |
| MF | 4.87e-05 | Enr | 2 | 1 | 0 | Yes | GO:0140657 | ATP-dependent activity |
| MF | 5.98e-05 | Enr | 3 | 1 | 0 | No | GO:0140296 | general transcription initiation factor binding |
| MF | 1.15e-04 | Enr | 2 | 1 | 0 | No | GO:0017111 | ribonucleoside triphosphate phosphatase activity |
| CC | 1.62e-04 | Enr | 4* | 1 | 0 | No | GO:0043232 | intracellular membraneless organelle |
| CC | 1.62e-04 | Enr | 4 | 1 | 0 | No | GO:0043228 | membraneless organelle |
| BP | 1.92e-04 | Enr | 1 | 1 | 0 | No | GO:0045596 | negative regulation of cell differentiation |
| MF | 1.92e-04 | Enr | 4 | 1 | 0 | No | GO:0001025 | RNA polymerase III general transcription initiation factor binding |
| BP | 1.92e-04 | Enr | 1* | 1 | 0 | No | GO:0010721 | negative regulation of cell development |
| MF | 1.92e-04 | Enr | 4* | 1 | 0 | No | GO:0001156 | TFIIIC-class transcription factor complex binding |
| BP | 1.92e-04 | Enr | 1* | 1 | 0 | No | GO:0075297 | negative regulation of ascospore formation |
