## Supplementary Files 1 to 5 for "*GOParsimony*: parsimonious saltation across the Gene Ontology digraph for efficient functional annotation": Suppl-File-4-Example-4-human-cancer-IDPs.pdf

#### Suppl. File 4: Graphical GOParsimony output for Example 4, the human cancer IDPs

**Figure A: Complete precedence graph (precedence set <=3).** The GO terms are labelled in the upper right corner with their precedence levels and are colour-graded with a blueness scale. The GO-term boxes also contain the term definition, the corrected P-value (Q-value if submitted to the Benjamini-Hochberg procedure), and the term type in the lower right corner (BP for biological process, CC for cellular component, MF for molecular function). They are connected with dotted arrows if there are only intervening non-significant GO terms. GOslim terms are outlined in magenta, and the user-specified highlight list in green. Here, the highlights come from default labelling of non-redundant GO terms with the Revigo website.

**Figure B: Concise precedence graph (precedence set <=3).** It is the same as in Figure A, except that only terms with precedence levels ≤ the level user-specified are included (here, the default = 5 is used).

**Figure C: 2D Heatmap for inferring functional modules. Proteins**/genes are clustered on the y-axis using a Jaccard distance calculated from the common GO terms. GO terms are clustered on the x-axis using a Jaccard distance calculated from the common proteins/genes. GO term names are colour-coded according to type (brown = CC, yellow = BP, green = MF). Cells are colour-coded for precedence as in the precedence graphs, and also labelled with the precedence level.

**Figure D: Comparison graph bar-chart indicating GO terms added as children or parents for higher precedence level thresholds.** Precedence level thresholds up to the precedence level specified in the GOParsimony run are shown (here, set ≤3).

**Figure E: Comparison graph Venn diagram.** The overlaps in the Precedence=1, default Revigo and GOslims term sets are shown. The table lists the percentage coverage of significant terms and the average –log(P-value) (i.e., corrected P-value).

**Figure F: Coloured GO term table.** It is colour-coded to correspond to the precedence graphs.

CONCISE PRECEDENCE GRAPHS (Prec <= 3)

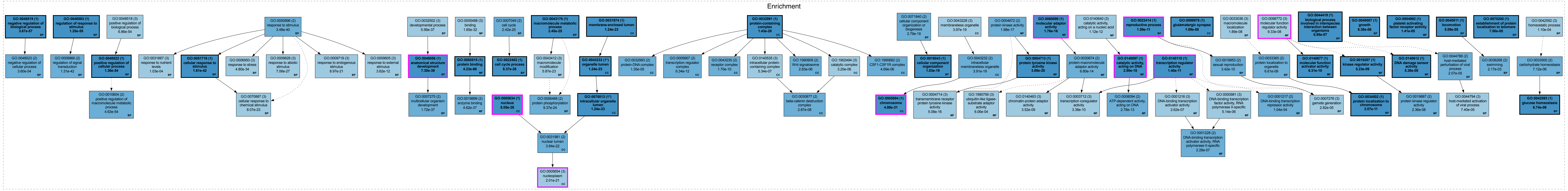

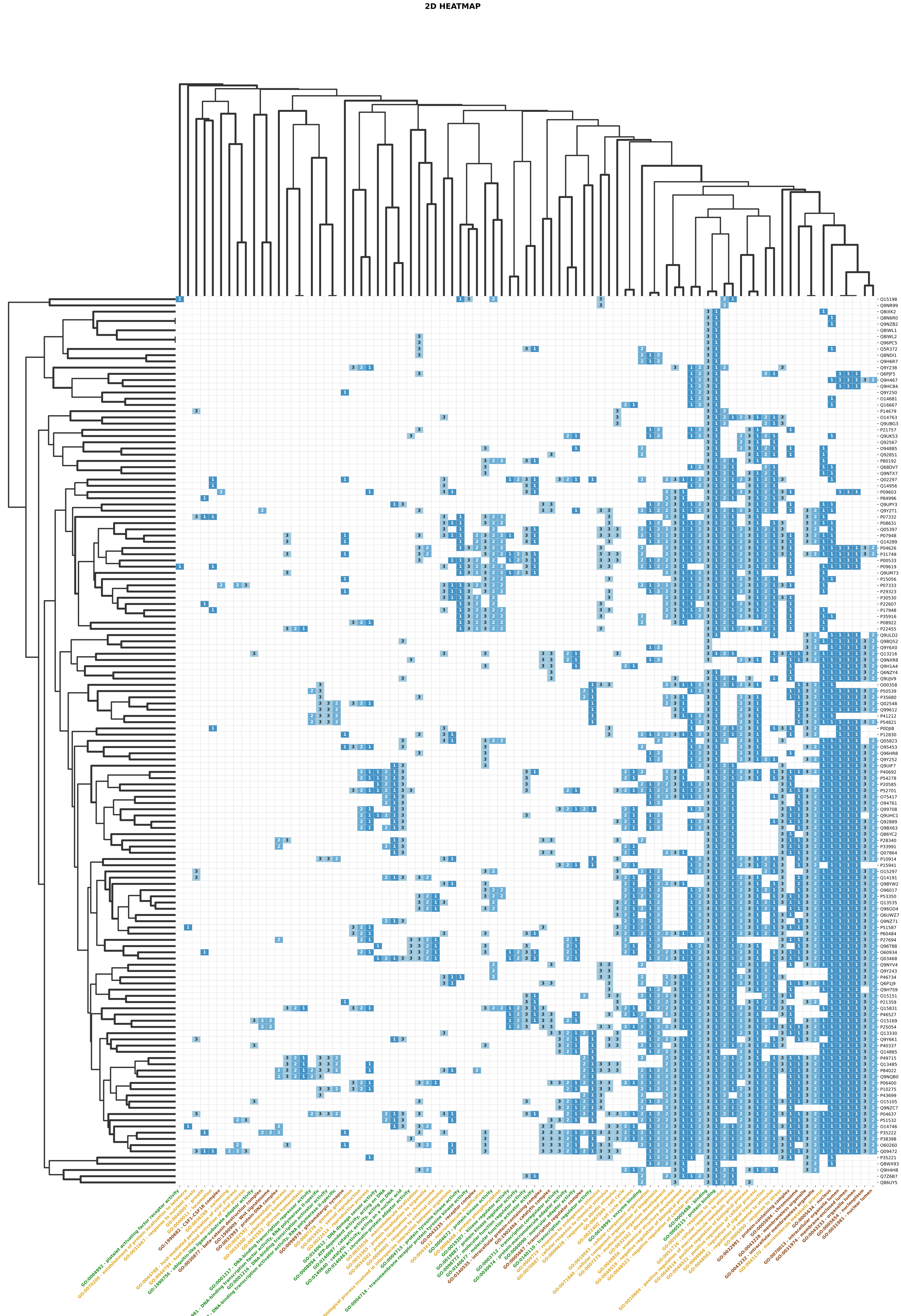
2D HEATMAP
Hierarchical Clustering of Shared GO Terms (Jaccard distance) [CC brown, BP yellow, MF green]

### COMPARISON GRAPHS

Functional Relationship of Added Terms relative to Precedence=1 Core

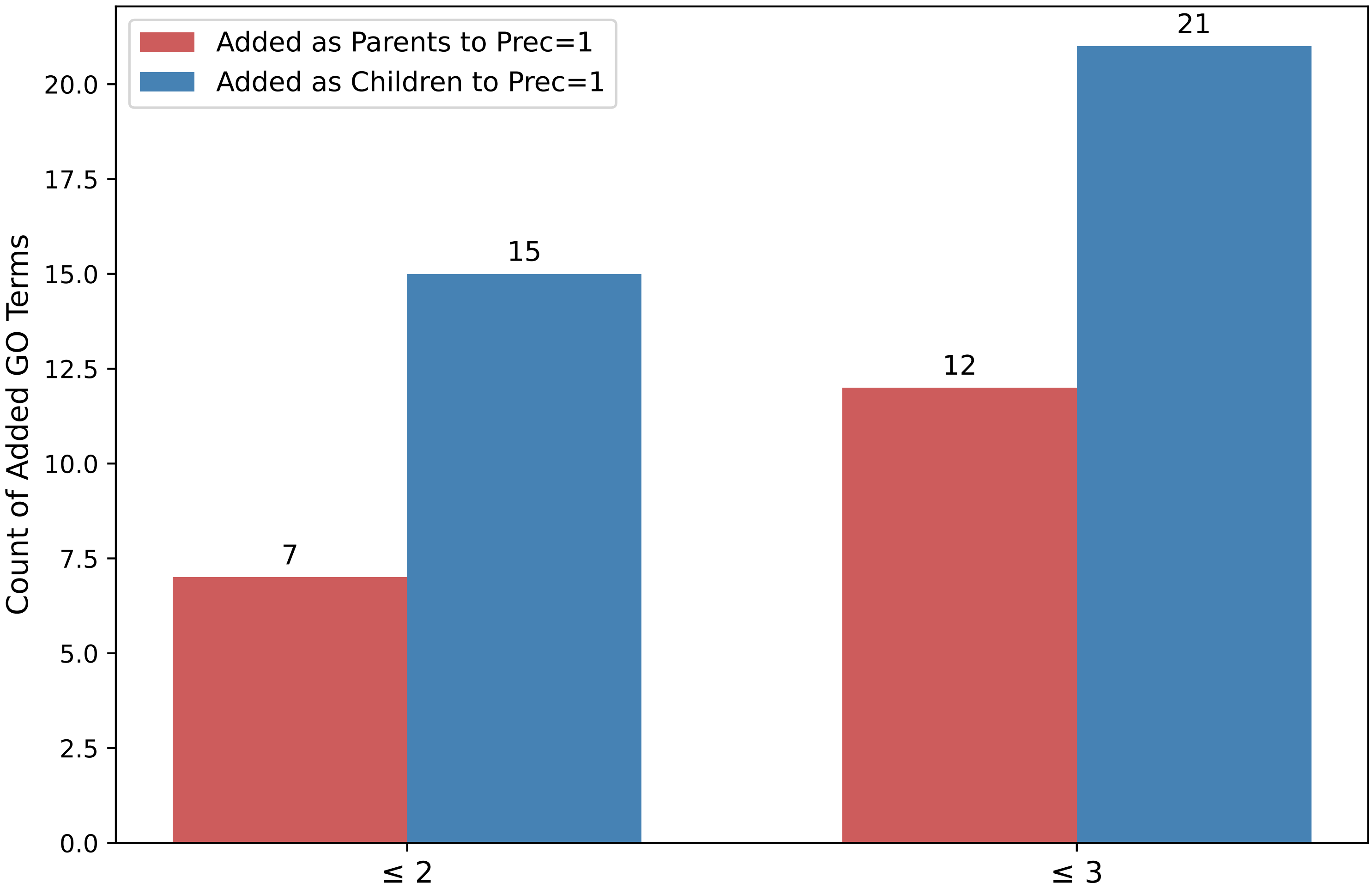

Overlap of Significant GO Terms

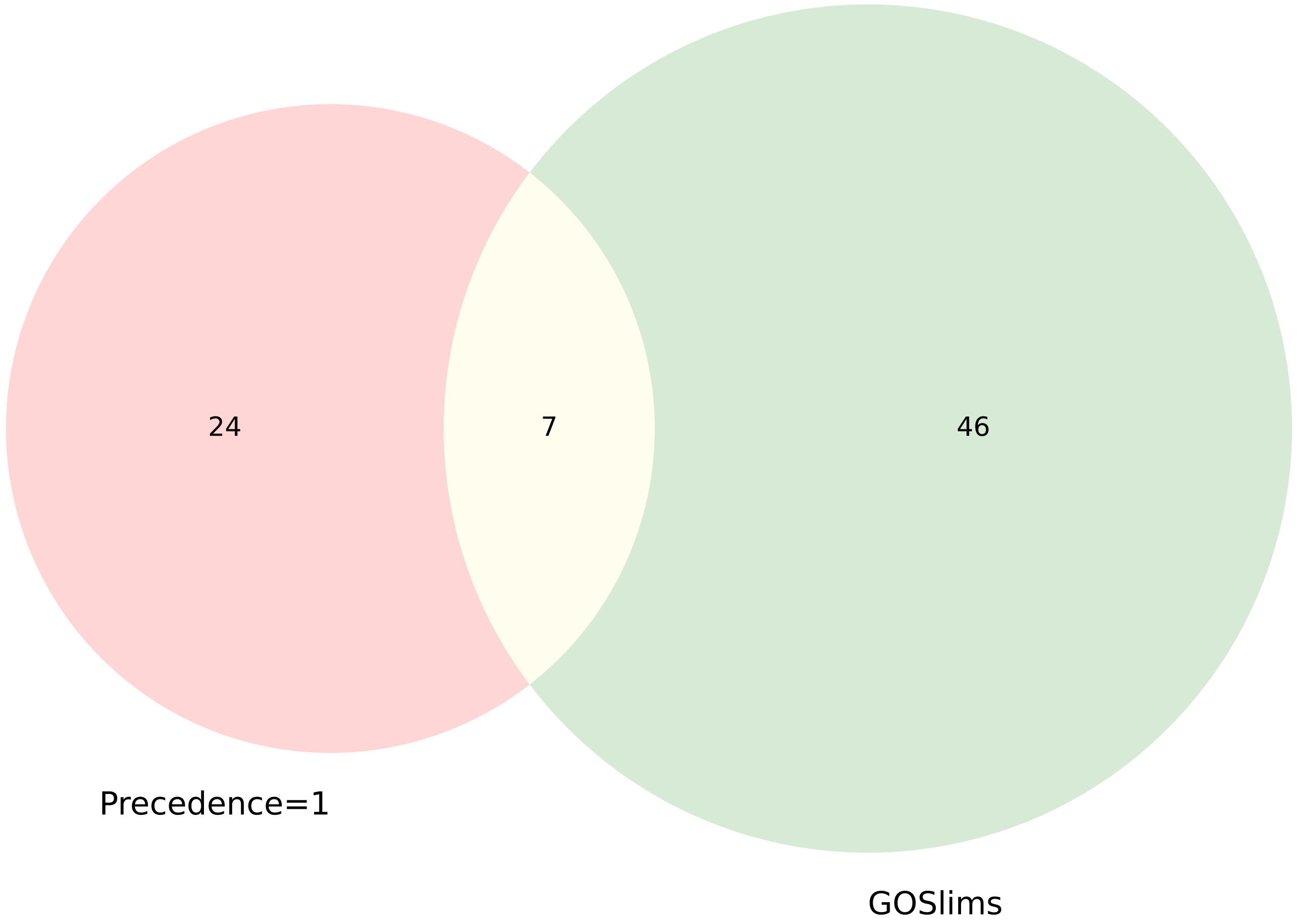


| Set Name | % Coverage (of the significant terms) | Average -log10(P-value) |
|---|---|---|
| **Precedence=1** | 1.8% | 21.64 |
| **GOSlims** | 3.1% | 10.64 |

### GOTERM TABLE (Prec <= 3)

| Namespace | P-Value | Enr/Dep | Prec | Reg_Prec | Reg_ClstID | Slim | GOTerm | Description |
|---|---|---|---|---|---|---|---|---|
| MF | 4.23e-66 | Enr | 1 | 1 | 0 | No | GO:0005515 | protein binding |
| BP | 3.67e-57 | Enr | 1 | 1 | 0 | No | GO:0048519 | negative regulation of biological process |
| BP | 1.25e-55 | Enr | 1 | 1 | 49 | No | GO:0048583 | regulation of response to stimulus |
| BP | 1.36e-54 | Enr | 1 | 1 | 50 | No | GO:0048522 | positive regulation of cellular process |
| BP | 3.60e-54 | Enr | 2 | 1 | 50 | No | GO:0048523 | negative regulation of cellular process |
| BP | 4.63e-54 | Enr | 2 | 1 | 51 | No | GO:0010604 | positive regulation of macromolecule metabolic process |
| BP | 6.86e-54 | Enr | 3 | 1 | 0 | No | GO:0048518 | positive regulation of biological process |
| BP | 1.31e-42 | Enr | 2 | 1 | 55 | No | GO:0009966 | regulation of signal transduction |
| BP | 1.61e-42 | Enr | 1 | 1 | 0 | No | GO:0051716 | cellular response to stimulus |
| BP | 3.48e-40 | Enr | 2 | 2 | 49 | No | GO:0050896 | response to stimulus |
| BP | 7.30e-39 | Enr | 1 | 1 | 0 | Yes | GO:0048856 | anatomical structure development |
| BP | 1.72e-37 | Enr | 2 | 1 | 63 | No | GO:0007275 | multicellular organism development |
| MF | 4.62e-37 | Enr | 2 | 1 | 0 | No | GO:0019899 | enzyme binding |
| BP | 5.56e-37 | Enr | 3 | 1 | 64 | No | GO:0032502 | developmental process |
| BP | 4.80e-34 | Enr | 3 | 1 | 70 | No | GO:0006950 | response to stress |
| MF | 1.65e-32 | Enr | 3 | 1 | 0 | No | GO:0005488 | binding |
| CC | 4.88e-31 | Enr | 1 | 1 | 0 | Yes | GO:0005694 | chromosome |
| BP | 7.06e-27 | Enr | 3 | 1 | 0 | No | GO:0009628 | response to abiotic stimulus |
| CC | 8.05e-26 | Enr | 1 | 1 | 0 | Yes | GO:0005634 | nucleus |
| BP | 8.37e-26 | Enr | 1 | 2 | 83 | No | GO:0022402 | cell cycle process |
| BP | 2.42e-25 | Enr | 2 | 2 | 75 | No | GO:0007049 | cell cycle |
| BP | 2.45e-25 | Enr | 1 | 2 | 51 | No | GO:0043170 | macromolecule metabolic process |
| BP | 9.37e-24 | Enr | 2 | 2 | 93 | No | GO:0006468 | protein phosphorylation |
| CC | 1.24e-23 | Enr | 1* | 1 | 0 | No | GO:0070013 | intracellular organelle lumen |
| CC | 1.24e-23 | Enr | 1* | 1 | 0 | No | GO:0043233 | organelle lumen |
| CC | 1.24e-23 | Enr | 1 | 1 | 0 | No | GO:0031974 | membrane-enclosed lumen |
| BP | 5.87e-23 | Enr | 3 | 1 | 0 | No | GO:0043412 | macromolecule modification |
| BP | 8.07e-23 | Enr | 3 | 1 | 0 | No | GO:0070887 | cellular response to chemical stimulus |
| CC | 3.84e-22 | Enr | 2 | 1 | 0 | No | GO:0031981 | nuclear lumen |
| CC | 2.01e-21 | Enr | 3 | 1 | 0 | Yes | GO:0005654 | nucleoplasm |
| BP | 8.97e-21 | Enr | 3 | 1 | 0 | No | GO:0009719 | response to endogenous stimulus |
| CC | 1.43e-20 | Enr | 1 | 1 | 0 | No | GO:0032991 | protein-containing complex |
| MF | 3.89e-20 | Enr | 1 | 1 | 0 | No | GO:0004713 | protein tyrosine kinase activity |
| BP | 1.03e-19 | Enr | 1 | 2 | 73 | No | GO:0016043 | cellular component organization |
| BP | 2.79e-19 | Enr | 2 | 1 | 0 | No | GO:0071840 | cellular component organization or biogenesis |
| CC | 3.91e-19 | Enr | 2 | 1 | 0 | No | GO:0043232 | intracellular membraneless organelle |
| CC | 3.97e-19 | Enr | 3 | 1 | 0 | No | GO:0043228 | membraneless organelle |
| MF | 2.98e-18 | Enr | 1 | 1 | 0 | Yes | GO:0140097 | catalytic activity, acting on DNA |
| MF | 1.98e-17 | Enr | 2 | 1 | 0 | No | GO:0004672 | protein kinase activity |
| MF | 1.76e-16 | Enr | 1 | 1 | 0 | Yes | GO:0060090 | molecular adaptor activity |
| MF | 5.08e-16 | Enr | 3 | 1 | 0 | No | GO:0004714 | transmembrane receptor protein tyrosine kinase activity |
| MF | 6.80e-14 | Enr | 2 | 1 | 0 | No | GO:0030674 | protein-macromolecule adaptor activity |
| MF | 2.78e-13 | Enr | 2 | 1 | 0 | No | GO:0008094 | ATP-dependent activity, acting on DNA |
| MF | 1.12e-12 | Enr | 3 | 1 | 0 | No | GO:0140640 | catalytic activity, acting on a nucleic acid |
| BP | 3.62e-12 | Enr | 3 | 2 | 176 | No | GO:0009605 | response to external stimulus |
| CC | 6.34e-12 | Enr | 2 | 1 | 0 | No | GO:0005667 | transcription regulator complex |
| MF | 1.40e-11 | Enr | 1 | 1 | 0 | Yes | GO:0140110 | transcription regulator activity |
| BP | 1.59e-11 | Enr | 1 | 1 | 0 | Yes | GO:0022414 | reproductive process |
| BP | 2.07e-11 | Enr | 1 | 1 | 0 | No | GO:0034502 | protein localization to chromosome |
| CC | 1.70e-10 | Enr | 2 | 1 | 0 | No | GO:0043235 | receptor complex |
| MF | 3.38e-10 | Enr | 3 | 1 | 0 | No | GO:0003712 | transcription coregulator activity |
| BP | 3.42e-10 | Enr | 2 | 1 | 0 | No | GO:0019953 | sexual reproduction |
| MF | 6.31e-10 | Enr | 1 | 1 | 0 | No | GO:0140677 | molecular function activator activity |
| MF | 3.52e-09 | Enr | 3 | 1 | 0 | No | GO:0140463 | chromatin-protein adaptor activity |
| MF | 5.23e-09 | Enr | 1 | 1 | 0 | No | GO:0019207 | kinase regulator activity |
| BP | 6.61e-09 | Enr | 2 | 1 | 0 | No | GO:0033365 | protein localization to organelle |
| CC | 1.09e-08 | Enr | 1 | 1 | 0 | No | GO:0098978 | glutamatergic synapse |
| BP | 1.89e-08 | Enr | 3 | 1 | 0 | No | GO:0033036 | macromolecule localization |
| MF | 2.36e-08 | Enr | 2 | 1 | 0 | No | GO:0019887 | protein kinase regulator activity |
| CC | 2.87e-08 | Enr | 2 | 1 | 0 | No | GO:0030877 | beta-catenin destruction complex |
| MF | 5.38e-08 | Enr | 1 | 1 | 0 | No | GO:0140612 | DNA damage sensor activity |
| MF | 9.33e-08 | Enr | 3 | 1 | 0 | Yes | GO:0098772 | molecular function regulator activity |
| MF | 2.28e-07 | Enr | 2 | 1 | 0 | No | GO:0001228 | DNA-binding transcription activator activity, RNA polymerase II-specific |
| MF | 2.62e-07 | Enr | 3 | 1 | 0 | No | GO:0001216 | DNA-binding transcription activator activity |
| CC | 5.34e-07 | Enr | 3 | 1 | 0 | No | GO:0140535 | intracellular protein-containing complex |
| BP | 6.95e-07 | Enr | 1 | 1 | 0 | No | GO:0044419 | biological process involved in interspecies interaction between organisms |
| CC | 2.83e-06 | Enr | 2 | 1 | 0 | No | GO:1990909 | Wnt signalosome |
| CC | 3.20e-06 | Enr | 3 | 1 | 0 | No | GO:1902494 | catalytic complex |
| CC | 4.69e-06 | Enr | 2 | 1 | 0 | No | GO:1990682 | CSF1-CSF1R complex |
| MF | 5.14e-06 | Enr | 3 | 1 | 0 | No | GO:0000981 | DNA-binding transcription factor activity, RNA polymerase II-specific |
| BP | 6.38e-06 | Enr | 1 | 2 | 126 | No | GO:0040007 | growth |
| BP | 6.74e-06 | Enr | 1 | 1 | 0 | No | GO:0042593 | glucose homeostasis |
| BP | 7.12e-06 | Enr | 2 | 1 | 0 | No | GO:0033500 | carbohydrate homeostasis |
| CC | 1.35e-05 | Enr | 2 | 1 | 0 | No | GO:0032993 | protein-DNA complex |
| MF | 1.41e-05 | Enr | 1 | 1 | 0 | No | GO:0004992 | platelet activating factor receptor activity |
| BP | 2.07e-05 | Enr | 2 | 1 | 0 | No | GO:0044788 | host-mediated perturbation of viral process |
| BP | 2.82e-05 | Enr | 3 | 1 | 0 | No | GO:0007276 | gamete generation |
| BP | 5.09e-05 | Enr | 1 | 2 | 110 | No | GO:0040011 | locomotion |
| BP | 7.00e-05 | Enr | 1 | 1 | 0 | No | GO:0070200 | establishment of protein localization to telomere |
| BP | 7.40e-05 | Enr | 3 | 1 | 0 | No | GO:0044794 | host-mediated activation of viral process |
| BP | 1.03e-04 | Enr | 3 | 1 | 0 | No | GO:0031667 | response to nutrient levels |
| MF | 1.04e-04 | Enr | 2 | 1 | 0 | No | GO:0001217 | DNA-binding transcription repressor activity |
| BP | 1.10e-04 | Enr | 3 | 1 | 0 | No | GO:0042592 | homeostatic process |
| MF | 6.06e-04 | Enr | 3 | 1 | 0 | No | GO:1990756 | ubiquitin-like ligase-substrate adaptor activity |
| BP | 2.17e-03 | Enr | 2 | 1 | 0 | No | GO:0036268 | swimming |
