## Supplementary Files 1 to 5 for "*GOParsimony*: parsimonious saltation across the Gene Ontology digraph for efficient functional annotation": Suppl-File-5-Example-5-human-lymphoblast-set.pdf

#### Suppl. File 5: Graphical GOParsimony output for Example 5, the human lymphoblast gene set, upregulated after treatment with ionizing radiation

**Figure A: Complete precedence graph (precedence set <=3).** The GO terms are labelled in the upper right corner with their precedence levels and are colour-graded with a blueness scale. The GO-term boxes also contain the term definition, the corrected P-value (Q-value if submitted to the Benjamini-Hochberg procedure), and the term type in the lower right corner (BP for biological process, CC for cellular component, MF for molecular function). They are connected with dotted arrows if there are only intervening non-significant GO terms. GOslim terms are outlined in magenta, and the user-specified highlight list in green. Here, the highlights come from default labelling of non-redundant GO terms with the Revigo website.

**Figure B: Concise precedence graph (precedence set <=3).** It is the same as in Figure A, except that only terms with precedence levels ≤ the level user-specified are included (here, the default = 5 is used).

**Figure C: 2D Heatmap for inferring functional modules. Proteins**/genes are clustered on the y-axis using a Jaccard distance calculated from the common GO terms. GO terms are clustered on the x-axis using a Jaccard distance calculated from the common proteins/genes. GO term names are colour-coded according to type (brown = CC, yellow = BP, green = MF). Cells are colour-coded for precedence as in the precedence graphs, and also labelled with the precedence level.

**Figure D: Comparison graph bar-chart indicating GO terms added as children or parents for higher precedence level thresholds.** Precedence level thresholds up to the precedence level specified in the GOParsimony run are shown (here, set ≤3).

**Figure E: Comparison graph Venn diagram.** The overlaps in the Precedence=1, default Revigo and GOslims term sets are shown. The table lists the percentage coverage of significant terms and the average –log(P-value) (i.e., corrected P-value).

**Figure F: Coloured GO term table.** It is colour-coded to correspond to the precedence graphs.

CONCISE PRECEDENCE GRAPHS (Prec <= 3)

Enrichment

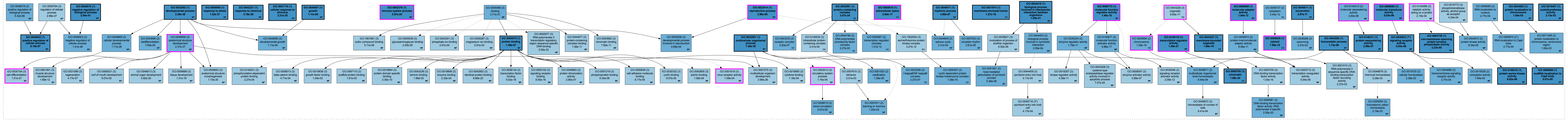

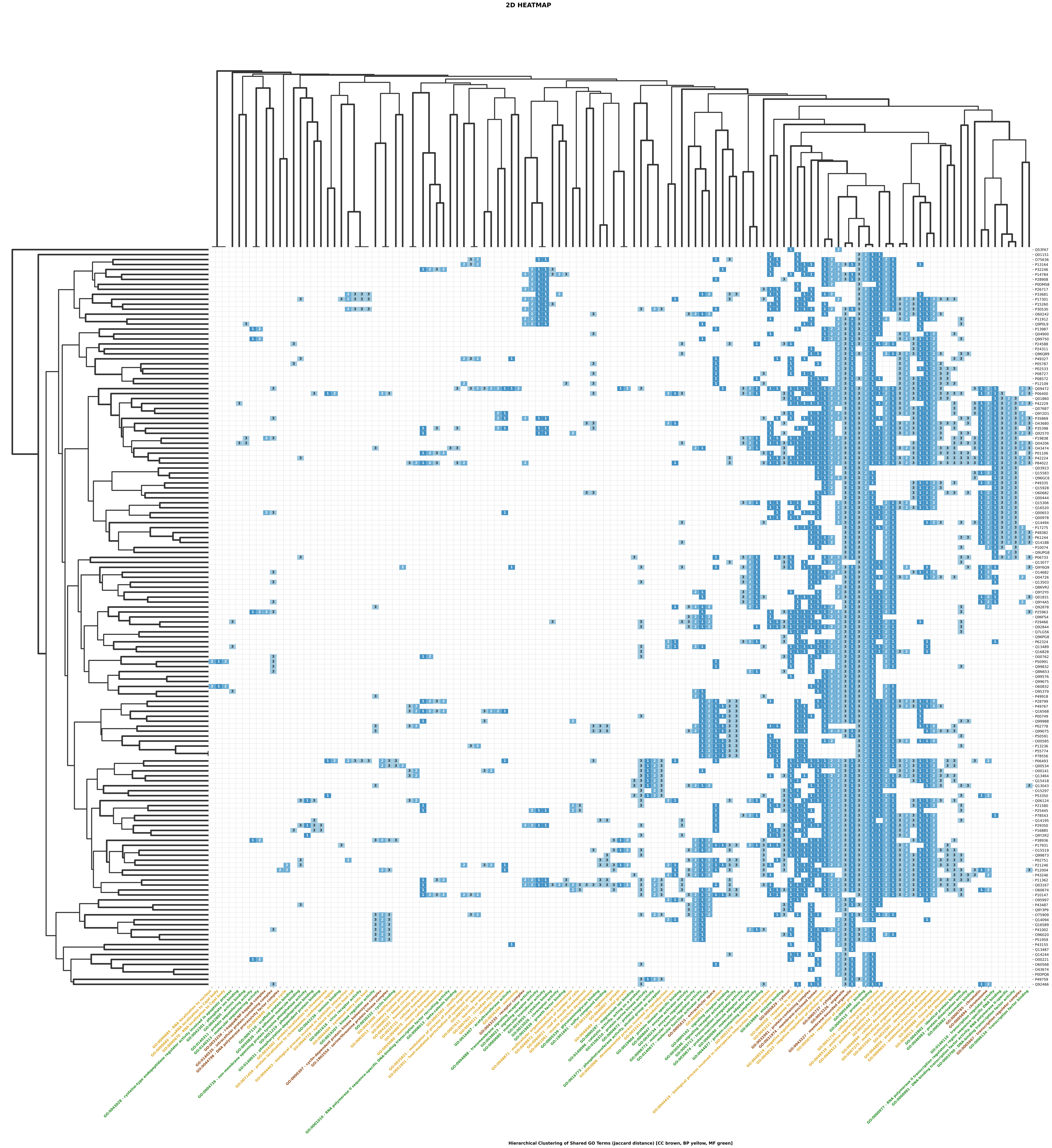
2D HEATMAP
Hierarchical Clustering of Shared GO Terms (Jaccard distance) [CC brown, BP yellow, MF green]

### COMPARISON GRAPHS

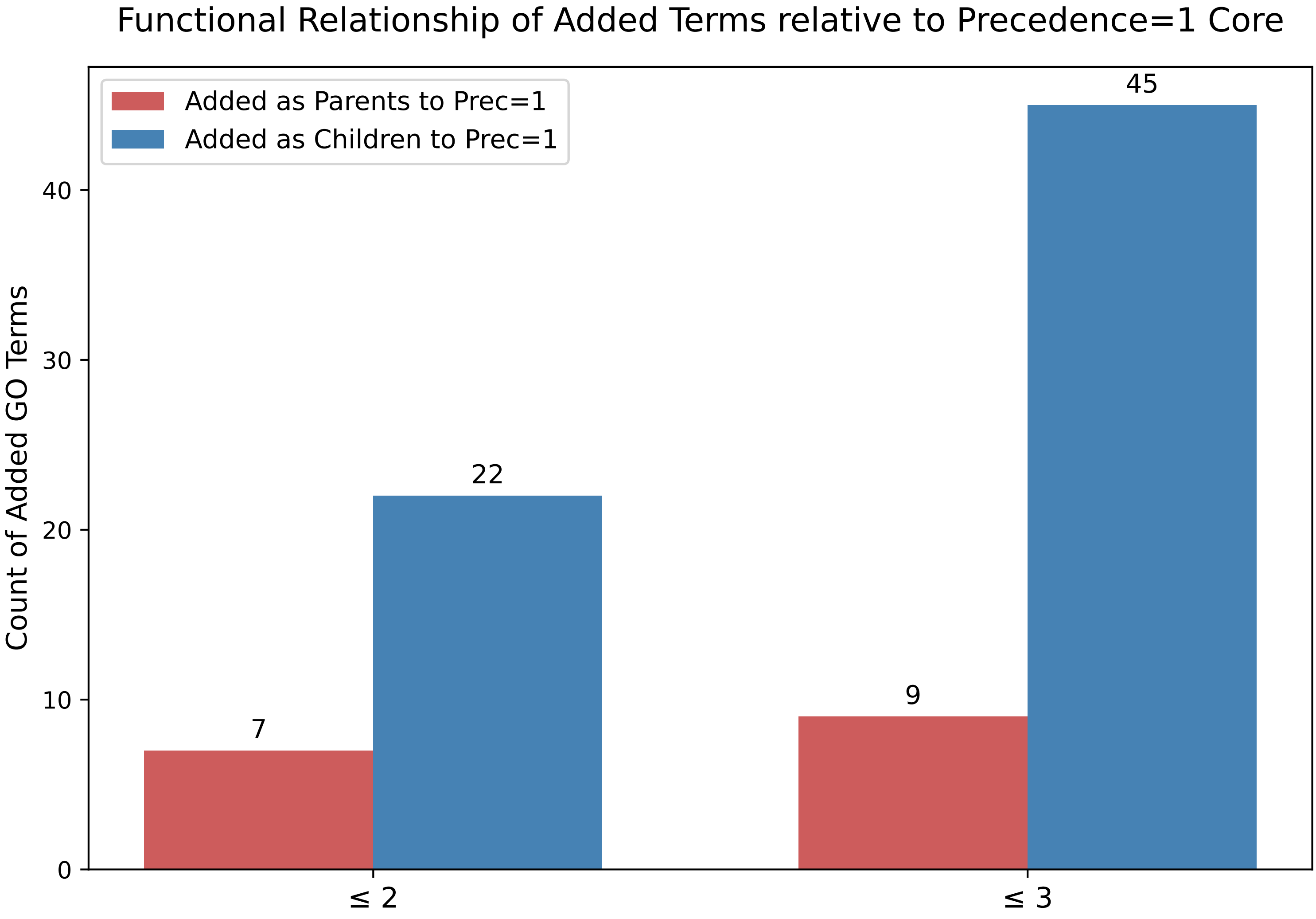

Overlap of Significant GO Terms

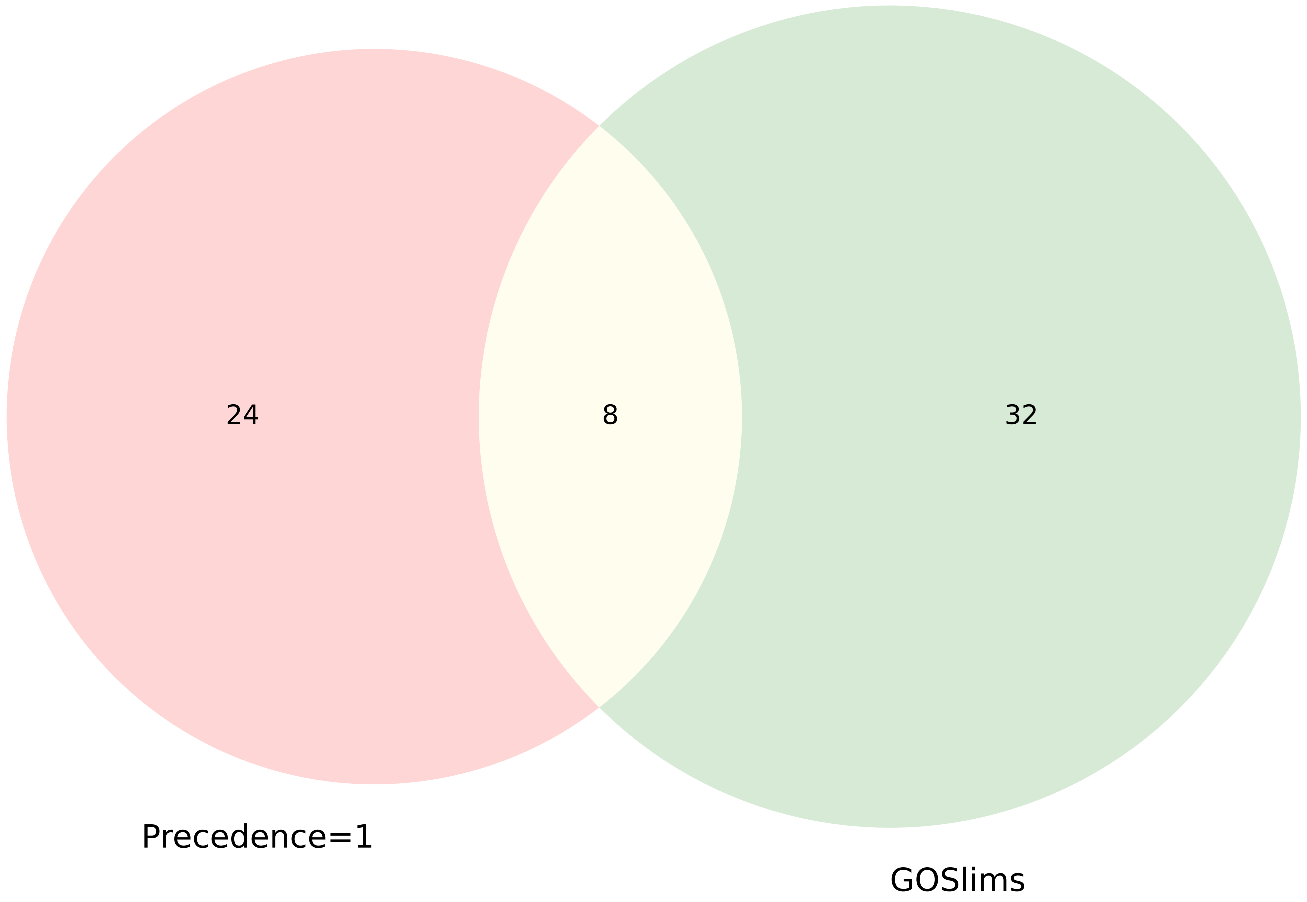


| Set Name | % Coverage (of the significant terms) | Average -log10(P-value) |
|---|---|---|
| **Precedence=1** | 2.2% | 19.76 |
| **GOSlims** | 2.7% | 11.39 |

### GOTERM TABLE (Prec <= 3)

| Namespace | P-Value | Enr/Dep | Prec | Reg_Prec | Reg_ClstID | Slim | GOTerm | Description |
|---|---|---|---|---|---|---|---|---|
| MF | 1.49e-67 | Enr | 1 | 1 | 0 | No | GO:0005515 | protein binding |
| BP | 6.19e-67 | Enr | 1 | 1 | 59 | No | GO:0048522 | positive regulation of cellular process |
| BP | 5.32e-66 | Enr | 2 | 1 | 0 | No | GO:0048518 | positive regulation of biological process |
| BP | 2.54e-51 | Enr | 1 | 1 | 0 | No | GO:0048519 | negative regulation of biological process |
| BP | 1.41e-48 | Enr | 2 | 1 | 59 | No | GO:0048523 | negative regulation of cellular process |
| BP | 2.99e-47 | Enr | 3 | 1 | 59 | No | GO:0050794 | regulation of cellular process |
| BP | 2.30e-38 | Enr | 1 | 1 | 69 | No | GO:0032502 | developmental process |
| BP | 1.32e-37 | Enr | 1 | 1 | 70 | No | GO:0006950 | response to stress |
| BP | 2.37e-37 | Enr | 2 | 1 | 0 | Yes | GO:0048856 | anatomical structure development |
| BP | 3.16e-36 | Enr | 1 | 1 | 0 | No | GO:0042221 | response to chemical |
| BP | 2.21e-35 | Enr | 1 | 1 | 0 | No | GO:0051716 | cellular response to stimulus |
| MF | 3.71e-31 | Enr | 2 | 1 | 0 | No | GO:0005488 | binding |
| BP | 1.38e-30 | Enr | 1 | 2 | 67 | No | GO:0032501 | multicellular organismal process |
| BP | 1.71e-28 | Enr | 2 | 1 | 0 | No | GO:0048869 | cellular developmental process |
| BP | 1.51e-27 | Enr | 3 | 1 | 87 | Yes | GO:0030154 | cell differentiation |
| BP | 2.96e-26 | Enr | 3 | 1 | 90 | No | GO:0007275 | multicellular organism development |
| CC | 1.58e-24 | Enr | 1 | 1 | 0 | No | GO:0000785 | chromatin |
| MF | 2.32e-24 | Enr | 3 | 1 | 0 | No | GO:0019899 | enzyme binding |
| BP | 2.57e-24 | Enr | 1 | 2 | 77 | Yes | GO:0002376 | immune system process |
| CC | 3.57e-24 | Enr | 1 | 1 | 0 | No | GO:0032991 | protein-containing complex |
| MF | 8.89e-23 | Enr | 3 | 1 | 0 | No | GO:0042802 | identical protein binding |
| MF | 2.65e-22 | Enr | 3 | 1 | 0 | No | GO:0000977 | RNA polymerase II transcription regulatory region sequence-specific DNA binding |
| BP | 1.05e-21 | Enr | 1 | 1 | 0 | No | GO:0044419 | biological process involved in interspecies interaction between organisms |
| MF | 1.36e-21 | Enr | 1 | 1 | 0 | Yes | GO:0140110 | transcription regulator activity |
| BP | 1.73e-20 | Enr | 3 | 1 | 106 | No | GO:0009653 | anatomical structure morphogenesis |
| MF | 2.50e-20 | Enr | 2 | 1 | 0 | No | GO:0000981 | DNA-binding transcription factor activity, RNA polymerase II-specific |
| BP | 4.82e-20 | Enr | 3 | 1 | 0 | No | GO:0048513 | animal organ development |
| CC | 1.29e-19 | Enr | 2 | 1 | 0 | Yes | GO:0005694 | chromosome |
| MF | 1.40e-18 | Enr | 1 | 1 | 0 | Yes | GO:0098772 | molecular function regulator activity |
| CC | 1.66e-18 | Enr | 1 | 1 | 0 | No | GO:0043227 | membrane-bounded organelle |
| MF | 5.02e-18 | Enr | 3 | 1 | 0 | No | GO:0008134 | transcription factor binding |
| CC | 2.94e-17 | Enr | 1 | 1 | 0 | Yes | GO:0005615 | extracellular space |
| MF | 5.86e-17 | Enr | 2 | 1 | 0 | No | GO:0140677 | molecular function activator activity |
| CC | 9.80e-17 | Enr | 3 | 1 | 0 | Yes | GO:0043226 | organelle |
| MF | 1.03e-16 | Enr | 3 | 1 | 0 | No | GO:0003700 | DNA-binding transcription factor activity |
| CC | 1.27e-16 | Enr | 1 | 1 | 0 | No | GO:0031974 | membrane-enclosed lumen |
| BP | 1.31e-16 | Enr | 3 | 1 | 0 | No | GO:0009888 | tissue development |
| MF | 8.55e-16 | Enr | 3 | 1 | 0 | No | GO:0005102 | signaling receptor binding |
| CC | 1.28e-15 | Enr | 2 | 1 | 0 | No | GO:0000307 | cyclin-dependent protein kinase holoenzyme complex |
| CC | 7.59e-15 | Enr | 1 | 1 | 0 | Yes | GO:0005829 | cytosol |
| CC | 1.01e-12 | Enr | 2 | 1 | 0 | No | GO:0005667 | transcription regulator complex |
| MF | 1.64e-12 | Enr | 1 | 1 | 0 | Yes | GO:0060090 | molecular adaptor activity |
| BP | 1.69e-12 | Enr | 3 | 1 | 0 | No | GO:0061061 | muscle structure development |
| MF | 2.09e-12 | Enr | 3 | 1 | 0 | No | GO:0030546 | signaling receptor activator activity |
| CC | 3.27e-12 | Enr | 3 | 1 | 0 | No | GO:1902554 | serine/threonine protein kinase complex |
| CC | 3.43e-12 | Enr | 2 | 1 | 0 | No | GO:0005737 | cytoplasm |
| MF | 1.06e-11 | Enr | 3 | 1 | 0 | No | GO:0044877 | protein-containing complex binding |
| MF | 1.79e-11 | Enr | 2 | 1 | 0 | No | GO:0030234 | enzyme regulator activity |
| BP | 2.87e-11 | Enr | 1 | 2 | 91 | No | GO:0040011 | locomotion |
| MF | 4.36e-11 | Enr | 3 | 1 | 0 | No | GO:0019207 | kinase regulator activity |
| MF | 6.06e-11 | Enr | 2 | 1 | 0 | No | GO:0030674 | protein-macromolecule adaptor activity |
| MF | 7.80e-11 | Enr | 3 | 1 | 0 | No | GO:0003682 | chromatin binding |
| MF | 3.30e-10 | Enr | 3 | 1 | 0 | No | GO:0019904 | protein domain specific binding |
| BP | 5.36e-09 | Enr | 2 | 1 | 0 | No | GO:0051851 | host-mediated perturbation of symbiont process |
| MF | 5.54e-09 | Enr | 3 | 1 | 0 | No | GO:0003712 | transcription coregulator activity |
| BP | 6.52e-09 | Enr | 3 | 1 | 0 | No | GO:0035821 | modulation of process of another organism |
| CC | 2.27e-07 | Enr | 2 | 1 | 0 | No | GO:0033256 | I-kappaB/NF-kappaB complex |
| MF | 2.80e-07 | Enr | 3 | 1 | 0 | No | GO:0046983 | protein dimerization activity |
| CC | 2.83e-07 | Enr | 2 | 1 | 0 | No | GO:0043235 | receptor complex |
| MF | 2.99e-07 | Enr | 1 | 1 | 0 | No | GO:0140311 | protein sequestering activity |
| MF | 3.88e-07 | Enr | 3 | 1 | 0 | No | GO:0008047 | enzyme activator activity |
| BP | 4.17e-07 | Enr | 3 | 1 | 0 | No | GO:0031099 | regeneration |
| BP | 4.90e-07 | Enr | 1 | 1 | 0 | No | GO:0048511 | rhythmic process |
| BP | 1.73e-06 | Enr | 1 | 1 | 0 | No | GO:0042592 | homeostatic process |
| MF | 2.60e-06 | Enr | 2 | 1 | 0 | Yes | GO:0140313 | molecular sequestering activity |
| MF | 5.11e-06 | Enr | 3 | 1 | 0 | No | GO:1901681 | sulfur compound binding |
| MF | 6.62e-06 | Enr | 1 | 1 | 0 | No | GO:0106310 | protein serine kinase activity |
| MF | 8.03e-06 | Enr | 1 | 1 | 0 | Yes | GO:0060089 | molecular transducer activity |
| MF | 8.03e-06 | Enr | 1* | 1 | 0 | No | GO:0038023 | signaling receptor activity |
| MF | 9.34e-06 | Enr | 3 | 1 | 0 | No | GO:0051219 | phosphoprotein binding |
| BP | 1.59e-05 | Enr | 2 | 2 | 389 | No | GO:0001503 | ossification |
| MF | 2.00e-05 | Enr | 3 | 1 | 0 | No | GO:0005539 | glycosaminoglycan binding |
| MF | 2.23e-05 | Enr | 1 | 1 | 0 | No | GO:0004726 | non-membrane spanning protein tyrosine phosphatase activity |
| BP | 2.61e-05 | Enr | 2 | 1 | 0 | No | GO:0007623 | circadian rhythm |
| BP | 2.96e-05 | Enr | 1 | 1 | 0 | Yes | GO:0022414 | reproductive process |
| BP | 5.07e-05 | Enr | 2 | 1 | 0 | No | GO:0008015 | blood circulation |
| BP | 5.07e-05 | Enr | 1 | 1 | 0 | No | GO:0090666 | scaRNA localization to Cajal body |
| MF | 6.17e-05 | Enr | 3 | 1 | 0 | No | GO:0050839 | cell adhesion molecule binding |
| MF | 7.07e-05 | Enr | 3 | 1 | 0 | No | GO:0043028 | cysteine-type endopeptidase regulator activity involved in apoptotic process |
| BP | 7.60e-05 | Enr | 2 | 1 | 0 | No | GO:0043696 | dedifferentiation |
| MF | 8.34e-05 | Enr | 2 | 1 | 0 | No | GO:0004672 | protein kinase activity |
| BP | 9.07e-05 | Enr | 3 | 1 | 0 | No | GO:0060021 | roof of mouth development |
| MF | 9.07e-05 | Enr | 3 | 1 | 0 | No | GO:0030332 | cyclin binding |
| MF | 1.06e-04 | Enr | 3 | 1 | 0 | No | GO:0042805 | actinin binding |
| MF | 1.09e-04 | Enr | 3 | 1 | 0 | Yes | GO:0001618 | virus receptor activity |
| BP | 1.11e-04 | Enr | 1 | 2 | 156 | No | GO:0040007 | growth |
| BP | 1.11e-04 | Enr | 2 | 2 | 456 | No | GO:0048589 | developmental growth |
| MF | 1.16e-04 | Enr | 3 | 1 | 0 | No | GO:0019955 | cytokine binding |
| BP | 1.74e-04 | Enr | 3 | 1 | 0 | Yes | GO:0003013 | circulatory system process |
| MF | 2.18e-04 | Enr | 3 | 1 | 0 | Yes | GO:0140096 | catalytic activity, acting on a protein |
| MF | 2.27e-04 | Enr | 3 | 1 | 0 | No | GO:0140031 | phosphorylation-dependent protein binding |
| CC | 2.41e-04 | Enr | 3 | 1 | 0 | No | GO:0140535 | intracellular protein-containing complex |
| BP | 2.59e-04 | Enr | 2 | 1 | 0 | No | GO:0044403 | biological process involved in symbiotic interaction |
| BP | 2.77e-04 | Enr | 2* | 1 | 0 | No | GO:0090670 | RNA localization to Cajal body |
| BP | 2.77e-04 | Enr | 2 | 1 | 0 | No | GO:0090685 | RNA localization to nucleus |
| BP | 3.31e-04 | Enr | 2 | 1 | 0 | No | GO:0044849 | estrous cycle |
| MF | 3.77e-04 | Enr | 2 | 1 | 0 | No | GO:0004888 | transmembrane signaling receptor activity |
| MF | 3.91e-04 | Enr | 3 | 1 | 0 | No | GO:0042301 | phosphate ion binding |
| BP | 4.17e-04 | Enr | 3 | 1 | 0 | No | GO:0044409 | symbiont entry into host |
| BP | 4.17e-04 | Enr | 3* | 1 | 0 | No | GO:0046718 | symbiont entry into host cell |
| MF | 4.24e-04 | Enr | 3 | 1 | 0 | No | GO:0016773 | phosphotransferase activity, alcohol group as acceptor |
| BP | 4.68e-04 | Enr | 2 | 1 | 0 | No | GO:0003006 | developmental process involved in reproduction |
| MF | 4.71e-04 | Enr | 3 | 1 | 0 | No | GO:0008013 | beta-catenin binding |
| MF | 7.94e-04 | Enr | 2 | 1 | 0 | No | GO:0015026 | coreceptor activity |
| BP | 8.44e-04 | Enr | 2 | 1 | 0 | No | GO:0048871 | multicellular organismal-level homeostasis |
| BP | 9.41e-04 | Enr | 3 | 1 | 0 | No | GO:0048872 | homeostasis of number of cells |
| MF | 1.04e-03 | Enr | 3 | 1 | 0 | No | GO:0019838 | growth factor binding |
| BP | 1.05e-03 | Enr | 1 | 1 | 0 | No | GO:0034502 | protein localization to chromosome |
| BP | 1.25e-03 | Enr | 2 | 1 | 0 | No | GO:0007611 | learning or memory |
| BP | 1.60e-03 | Enr | 2 | 1 | 0 | No | GO:0071459 | protein localization to chromosome, centromeric region |
| MF | 1.61e-03 | Enr | 3 | 1 | 0 | No | GO:0097110 | scaffold protein binding |
| MF | 1.99e-03 | Enr | 3 | 1 | 0 | No | GO:0043236 | laminin binding |
| BP | 2.01e-03 | Enr | 3 | 2 | 760 | No | GO:0007610 | behavior |
| MF | 2.12e-03 | Enr | 1 | 1 | 0 | No | GO:0016407 | acetyltransferase activity |
| BP | 2.19e-03 | Enr | 2 | 1 | 0 | No | GO:0055080 | monoatomic cation homeostasis |
| CC | 2.27e-03 | Enr | 2 | 1 | 0 | No | GO:0044796 | DNA polymerase processivity factor complex |
| BP | 2.27e-03 | Enr | 2 | 1 | 0 | No | GO:0036268 | swimming |
| MF | 2.27e-03 | Enr | 3 | 1 | 0 | No | GO:0001010 | RNA polymerase II sequence-specific DNA-binding transcription factor recruiting activity |
| BP | 2.28e-03 | Enr | 3 | 1 | 0 | No | GO:0048878 | chemical homeostasis |
| BP | 2.38e-03 | Enr | 2 | 1 | 0 | No | GO:0019725 | cellular homeostasis |
| MF | 2.91e-03 | Enr | 3 | 1 | 0 | No | GO:0000287 | magnesium ion binding |
